# AnnoMiner: a new web-tool to integrate epigenetics, transcription factor occupancy and transcriptomics data to predict transcriptional regulators

**DOI:** 10.1101/2021.01.28.428613

**Authors:** Arno Meiler, Fabio Marchiano, Michaela Weikunat, Frank Schnorrer, Bianca H. Habermann

## Abstract

Gene expression regulation requires precise transcriptional programs, led by transcription factors in combination with epigenetic events. Recent advances in epigenomic and transcriptomic techniques provided insight into different gene regulation mechanisms. However, to date it remains challenging to understand how combinations of transcription factors together with epigenetic events control cell-type specific gene expression. We developed the AnnoMiner web-server and introduce an innovative and flexible way to annotate and integrate epigenetic, and transcription factor occupancy data. First, AnnoMiner annotates user-provided peaks with gene features. Second, AnnoMiner can integrate genome binding data from two different transcriptional regulators together with gene features. Third, AnnoMiner offers to explore the transcriptional deregulation of 10 genes nearby a user-provided peak. AnnoMiner’s fourth function performs transcription factor or histone mark enrichment analysis for user-provided gene lists by utilizing hundreds of public, high-quality datasets from ENCODE for the model organisms human, mouse, *Drosophila* and *C. elegans*. Thus, AnnoMiner can predict transcriptional regulators for a studied process without the strict need for chromatin data from the same process. We compared AnnoMiner to existing tools and experimentally validated several transcriptional regulators predicted by AnnoMiner to indeed contribute to muscle morphogenesis in *Drosophila*. AnnoMiner is freely available at http://chimborazo.ibdm.univ-mrs.fr/AnnoMiner/.

## Introduction

Transcriptional regulation is a highly complex process involving a combination of various molecular players and biochemical mechanisms, such as transcription factors (TFs), histone modifying enzymes (HMs), DNA methylases, as well as a structural reorganization of chromatin. Technical advances in analysing the interaction of proteins with DNA (ChIP-seq), to detect open or closed chromatin states (e.g. ATAC-seq, DNase-seq, FAIRE-seq), to detect hypermethylated CpG islands (bisulfite sequencing), or to map higher order chromosomal structural organisation (ChiaPET, Hi-C, 3C-seq) have revolutionized and significantly advanced our understanding of transcriptional regulation during the last decade (1, 2). Among other things, transcriptional enhancers were identified as crucial for regulating spatio-temporal gene expression programs by interacting with target gene promoters, often across large genomic distances (see (3–5) and references therein). Furthermore, the genome sequence in the chromatin is not a simple linear thread but organized in 3D, forming compartments, topologically associated domains (TADs) or chromatin loops that can bring distant elements in proximity, all of which can contribute to transcriptional regulation ((3, 6) and references therein). More recent evidence suggests even the presence of dual-action cis-regulatory modules (CRMs) that act as promoters, as well as distal enhancers (7, 8). All these findings suggest that transcriptional regulation is far more complex than initially anticipated and involves a collective effort of specific binding sites in the genome, a complex genome structure and the presence of various transcriptional regulators. Hence, we need a tool that can ideally integrate all this information to better understand and predict transcriptional regulation.

Techniques such as ChIP-seq, ATAC-seq, Hi-C seq and others involve an NGS (Next Generation Sequencing) step, resulting typically in paired-end reads of isolated chromosomal fragments. Therefore, the first steps after sequencing consist of read mapping, which is usually done using a software such as BowTie2 (9) followed by peak calling. There are a number of tools available for peak calling (reviewed e.g. in (10–13)), which include MACS2 (14) for standard peak calling, ChIPdiff (15), EpiCenter (16) or diffReps (17) for differential peak calling. The output of peak callers are the genomic coordinates of epigenetic marks or transcription factor binding sites under study. These coordinates are commonly stored in the Browser Extensible Data (BED) file format, a light-weight, standardized format to share genome coordinates.

The next step to biologically interpret genomic coordinates (also called peaks) is their genomic annotation, a process referred to as peak annotation or gene assignment. A number of peak annotation tools exist. Some of them combine ChIP-seq data analysis (including peak calling) and peak annotation, including the ChIP-Seq tools and web server ((18), web-based), Sole-Search (19), CIPHER (20), Nebula ((21), web-based, server no longer reachable), PeakAnalyser (22), BEDTools (23) or HOMER (24). Some of them are specific to peak annotation and visualization, such as ChIPseeker (25), UROPA (26), annoPeak ((27), web-based, server no longer reachable), ChIPseek ((28), web-based, server not responding), PAVIS ((29), web-based), Goldmine (30), GREAT ((31), web-based), or ChIPpeakAnno (32). While most of the peak annotators assign peaks to the closest TSS or genome feature, others consider up- and down-stream gene features, or provide the overlap with gene feature attributes (such as promoter, 5’UTR, 3’UTR, exon, intron) of the nearest gene (18, 21, 22, 25, 26, 29). The web-tool GREAT (31) offers peak annotation and gene assignment in larger genomic regions based on Gene Ontology (GO)-term similarity. What appears missing is a web-based, visual tool that helps experimental biologists to explore and integrate peak data with transcriptomics data in a flexible, user-centric and easy way. AnnoMiner is designed to close this gap.

An enormous community effort has been invoked in the past decade to collect, standardize and present annotated genomic data in form of the ENCODE (33–35), modENCODE (36) and modERN (37) resources, with the aim to make sense of the encyclopaedias of genomes. These initiatives have also made it possible to explore available genomic data further and integrate them with each other as well as with user-generated data. These standardized, high-quality data can be used to identify enriched transcription factor binding events in promoters of co-regulated genes, for example from a differential gene expression dataset (e.g. an RNA-seq dataset). This form of data integration predicts possible transcriptional regulators for biological processes under study, which is already widely used in the community. Enrichment analysis is commonly performed by testing for TF overrepresentation in the promoter regions of a user-provided gene list compared to a background list (e.g. considering the entire genome). Most available tools define the promoter region rigidly as a range of upstream and downstream base pairs from the annotated gene transcription starting site (TSS) and these parameters are kept fixed for all the TFs tested, without accounting for the differences between individual transcription factors. Following this approach, divers web-based tools have been developed, including VIPER (38), DoRothEA (39), BART (40), oPOSSUM (41), TFEA.ChIP (42), ChEA3 (43), EnrichR (44) or i-cisTarget (45). However, it would be advantageous to have a web-based, flexible and user-friendly solution for TF enrichment, considering promoter boundaries specific to each TF and working for the most widely used species (human, mouse, *Drosophila* and *C. elegans*).

Here we present AnnoMiner, a flexible, web-based, and user-friendly platform for peak annotation and integration, as well as TF and HM enrichment analysis. AnnoMiner allows users to annotate and integrate multiple genomic regions files with gene feature attributes and with transcriptomic data in an interactive and flexible way. AnnoMiner contains three distinct functions with different genomic peak annotation purposes in mind: first, *peak annotation*, which can be used to assign attributes of gene features to peaks, including user-defined upstream and downstream regions, TSS, 5’ and 3’UTRs, as well as the coding region; second, *peak integration* to search for overlapping binding events of two transcriptional regulators (e.g. two different TFs; two different HMs; or a TF in combination with a HM) with gene feature attributes; and third, *nearby genes annotation*, which helps to identify long range interaction effects on gene expression of a single genomic region. All three annotation features can optionally integrate user provided data, such as results from differential gene expression analysis, allowing to inspect for instance expression data and genomic peak data from multiple transcriptional regulators together. As a fourth function, AnnoMiner performs *TF (Transcription factor) and HM (Histone Mark) enrichment analysis*, using all high-quality filtered TF and HM data from ENCODE, modENCODE and modERN, which are stored in an internal database, and a user defined gene lists as input. AnnoMiner’s *TF enrichment function* offers the DynamicRanges option, which uses promoter regions specific to each TF based on pre-calculated binding densities for each individual TF. We tested the predictive power of AnnoMiner’s *TF enrichment function* and applied it to flight muscle development in *Drosophila* focusing on the process of myofibril morphogenesis (46). AnnoMiner predicted two potential transcriptional regulators, Trithorax-like (Trl) and the uncharacterized zinc-finger protein CG14655 to play a role during this process. For both, we provide experimental evidence for an essential function in myofibril morphogenesis in flight muscle. Thus, AnnoMiner correctly predicted a new role for Trl, as well as CG14655 as transcriptional regulators in a specific muscle type in *Drosophila*. Finally, we used AnnoMiner to predict direct transcriptional targets for the likewise enriched TFs Yorkie (Yki) and Scalloped (Sd) required for flight muscle growth (47).

## Methods

### The AnnoMiner database

The basic structure of the AnnoMiner web-server is shown in Supplementary Figure S1. In the back-end, all genomic data (retrieved from UCSC), data from ENCODE (33), modENCODE (36) and modERN (37), as well as user uploads are stored in the document-oriented, non-relational database MongoDB. This type of database structure was chosen for its flexibility and its speed. For each model organism, the latest genome assembly is stored. For human, mouse and *Drosophila*, we also store the second latest release. We use Java Servlets and Java Connectors for analysis, data upload, and connection to MongoDB, respectively. The *front-end* is based on Javascript and HTML, whereby the Annominer.js performs all Post calls to the Java servlets in the *back-end*. To follow the Findability, Accessibility, Interoperability and Reusability (FAIR) principle (48), documentation and source code of the tool are available on GitLab: https://gitlab.com/habermann_lab/annominer. All python scripts are modular and stored individually, allowing local usage for different purposes, and to populate the MongoDB database with ENCODE, modENCODE or modERN data. Java applets can equally be used to perform batch uploads, download genome data, define and compute DynamicRanges in a customizable way and finally, to reproduce our benchmark analysis.

### Genomic peak annotation functions

#### Peak annotation

For calculating the overlap between peaks and genes, we compare the coordinates of peaks with those of stored gene feature attributes. These include 5’ flanking regions, promoter region, transcription start site (TSS), 5’UTR, coding sequence (CDS, including exons and introns), 3’UTR, 3’ flanking regions, as well as strandedness of the gene feature. For integrating peaks with gene features, we first calculate the coverage of each peak by testing whether the peak coordinates fall within the coordinates of any attribute of a gene feature. If all isoforms of a gene are chosen, the overlap of the peak with the attributes of each individual isoform will be checked. For calculating the coverage, overlap events are calculated cumulatively and the sums of all overlaps will be divided by the length of the attribute. Depending on which attribute (target region) the user has chosen, only genes whose chosen attribute overlaps (with a minimum % or number of base pairs) with the peak will be displayed in the data table.

#### Peak integration

The *peak integration* function is based on the same algorithm as the *peak annotation*. However, only genes, which meet the overlap criteria of the user-selected target regions with peaks from both BED files will be displayed in the resulting data table.

#### Nearby genes annotation

For the *nearby genes annotation* function, the 5 neighbouring genes up- and down-stream, as well as the overlapping gene of a peak are extracted and displayed to the user. The user can choose whether the directionality of the gene with respect to the peak should be considered. If a gene overlaps with the peak, it will be shown in the *Overlap* box. Data on differential expression, which needs to be provided by the user in form of an annotation file, is displayed on the returned plot. The user can choose whether or not the directionality of the deregulation (up- or downregulated) is shown in the box plot. Depending on the user input, the *nearby genes annotation* function returns neighbouring genes of a single peak, or of many peaks.

### Transcription factor enrichment analysis

Transcription factor enrichment analysis in AnnoMiner uses publicly available TF ChIP-seq data from the ENCODE, modENCODE and modERN databases. In order to keep only high-quality data, we undertook a strict selection of available TF ChIP-seq datasets. First, we kept only experiments, for which data had already been pre-analysed following the ENCODE processing pipeline. Second, only experiments with replicates and optimal IDR peaks files (narrowPeak) were retained. Third, we manually inspected each narrowPeak file, evaluating the peak distribution around the gene’s TSS and discarded the ones with irregular peak distributions in relation to promoter regions of genes, as we considered those as outliers (Supplementary Figure S2 c, d; all outliers csv files are available in our gitlab repository at https://gitlab.com/habermann_lab/AnnoMiner-paper). The number of used TF ChIP-seq datasets for each of the model organisms and assemblies is available in Supplementary Tables S1, S2.

#### Over-representation analysis

For each TF ChIP-seq dataset stored in the AnnoMiner database, we first compute the contingency table of potential targets in both the uploaded gene list (signature gene-set) and in the background list of all genes of the chosen organism. We then apply a one-tailed hypergeometric test followed by Benjamini-Hochberg correction (49). Furthermore, we calculate the enrichment score as follows:

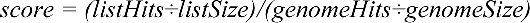

And a combined score based on the p-value and enrichment score according to (50):

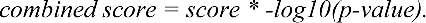

#### Promoter definition

AnnoMiner allows the user to fully customize the promoter region considered for TF enrichment analysis. The user can choose both the upstream and downstream borders to consider without constraints.

#### DynamicRanges option

The user can also choose dynamic ranges that we pre-calculated for each individual TF based on available ChIP-seq data stored in AnnoMiner. In brief, for each TF ChIP-seq experiment we computed the distances between its peaks and the genes in a range of 20,000 bp upstream of their TSSs. We then binned the closest genes by distances with a resolution of 50 bp. A noise level was computed, representing non-significant bindings, from 5,000 to 20,000 bp, as follows:

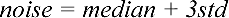

Next, we smoothened the binning using the simple moving average approach (SMA) with a sliding window of size=3. Intersecting the noise threshold with the SMA curve, we obtained the upstream TSS value. This value is unique for each TF and represents the distance range in which the TF is more often binding than is expected at random (see Supplementary Figure S2 e, f).

#### Choosing the peak - promoter overlap

The user can define the amount of overlap between a TF and a promoter in percentage or in base pairs (bp) to categorize a gene as a potential target.

### Histone mark enrichment analysis

The same function that is used for the *Transcription factor enrichment analysis* can be used for enriching histone marks for a gene list. In this case, a *DynamicRanges*-like option is not meaningful. Thus, the user has to manually define the promoter region (default: 2000bp upstream and 500bp downstream of a gene’s TSS).

### Automatic ID conversion

In order to easily allow comparisons between genome versions (ENSEMBL, RefSeq, UCSC, Genecode, Flybase and Wormbase) and not restrict the user to a specific set of gene identifiers, we developed a geneID converter. This function converts on the fly the user’s input gene IDs to match the selected resource, without any intervention from the user. The conversion is performed using the latest release of BioMart (51). BioMart data were downloaded and are stored locally for reasons of speed and are regularly updated.

### Benchmarking the TF enrichment function of AnnoMiner

#### Benchmarking dataset

Benchmarking data-sets for testing AnnoMiner’s enrichment function in the form of TF regulons were obtained from human single TF perturbation experiments (knock-downs, knockouts, over-expression and chemical inhibition) followed by RNA-seq from Gene Expression Omnibus (GEO (52), 160 experiments), manually curated and made available from Keenan et al. (43). We retained only experiments targeting TFs for which AnnoMiner is storing at least one high quality TF ChIP-seq dataset for the assembly GRCh38 (hg38), leading to a total of 75 RNA-seq experiments left for benchmarking (*benchmarkingSet*).

#### Benchmarking metrics

Precision-Recall (PR) and Receiver-Operating Characteristic (ROC) areas under the curves (AUC) for the *benchmarkingSet* were computed using the R package PRROC (53), following the benchmarking procedure proposed and applied by (43) and (54). In brief, for each TF, signature gene-sets were derived based on differentially expressed genes of the perturbed TF versus its control and used for TF enrichment analysis. For each dataset, the rank of the perturbed TF was assigned to the positive (foreground (*fg*)) class, while all other ranks of TFs retrieved for the dataset were assigned to the negative (background (*bg)*) class; using the PPROC package, precision-recall (PR) and Receiver-Operator Curves (ROC) using continuous interpolation were computed and areas under curve (AUCs) for both were calculated (PR-AUC and ROC-AUC). As the negative *bg* class greatly exceeded the positive *fg* class, we down-sampled the negative class to the same size as the positive class. We repeated this procedure 5000 times. We used the *approx* R function to linearly interpolate between all points from the 5000 ROC and the 5000 PR curves and calculated the mean ROC and PR AUCs values of all bootstrap runs. Furthermore, we analysed the cumulative distribution of the rank values for each TF across all experiments it was perturbed in, which is not expected to be uniform, unless the TF would rank randomly in all its associated datasets. We then performed the Anderson-Darling test using the *goftest* R package to test for significant deviations from this distribution, indicating a non-random distribution of the TF. Finally, we computed the percentage of perturbed TFs that were correctly ranked within the first percentile to ensure high ranking of the TF. All values are available in Table 1.

**Table 1:** performance values of AnnoMiner compared to other TF enrichment web-tools. AnnoMiner’s metrics are reported only for RefSeq and the parameters: upstream_tss: DynamicRange, downstream_tss:50bp and overlap:20bp.

| *Tool* | *Method* | *ROC*<br>*AUC* | *PR*<br>*AUC* | *%*<br>*recovered* | *ADtest* | *# of*<br>*datasets* |
| --- | --- | --- | --- | --- | --- | --- |
| ChEA3 | meanRank | 0.83 | 0.81 | 6.7 | 8.00E-06 | 75 |
|  | topRank | 0.82 | 0.81 | 9.3 | 8.00E-06 |  |
| AnnoMiner |  | 0.63 | 0.63 | 4.0 | 8.10E-05 |  |
| TFEA.ChIP |  | 0.63 | 0.60 | 0.0 | 7.90E-04 | 57 |
| AnnoMiner |  | 0.67 | 0.67 | 5.3 | 1.07E-05 |  |
| EnrichR | TRRUST_Transcription_Factors_2019 | 0.71 | 0.72 | 7.8 | 1.18E-05 | 51 |
| AnnoMiner |  | 0.63 | 0.65 | 5.9 | 5.57E-04 |  |
| **EnrichR** | ENCODE_TF_ChIP-seq_2015 | 0.68 | 0.68 | 4.6 | 3.59E-05 | 44 |
| **AnnoMiner** |  | 0.72 | 0.71 | 6.8 | 1.36E-05 |  |
| **EnrichR** | ChEA_2016 | 0.70 | 0.72 | 18.2 | 1.36E-05 | 44 |
| **AnnoMiner** |  | 0.67 | 0.68 | 4.6 | 5.12E-05 |  |
| **EnrichR** | ARCHS4_TFs_Coexp | 0.65 | 0.67 | 6.8 | 8.11E-06 | 74 |
| **AnnoMiner** |  | 0.63 | 0.64 | 4.1 | 6.05E-05 |  |
| **EnrichR** | ENCODE_and_ChEA_Consensus | 0.57 | 0.60 | 10.3 | 0.27 | 29 |
| **AnnoMiner** |  | 0.76 | 0.75 | 6.9 | 2.07E-05 |  |

### Benchmarking against existing tools

We compared the performance of AnnoMiner’s TF enrichment function with the performance of the following web-servers: TFEA.ChIP (42), ChEA3 (43) and EnrichR (44). The same dataset (*benchmarkingSet*) was used for all web-tools tested. Only datasets which could be effectively ranked by the web-tool under study were used. We used 57 signature gene-sets to test TFEA.ChIP and all the 75 signature-gene sets to test ChEA3. For EnrichR we evaluated the performance metrics on 5 different resources: ARCHS4, ChEA 2016, ENCODE 2015, ENCODE and ChEA Consensus and TRRUST 2019 using 74, 44, 44, 29 and 51 signature gene-sets, respectively.

### Epigenetic and RNA-seq data used

For analysing STAT3 in acute B-cell lymphomas, we used data from super-series GSE50724 (55). Only peaks which were significantly upregulated in ABC type B-cell lymphoma were selected for AnnoMiner’s *peak integration* (FDR cut-off <= 0.05; FC >= 1.25). We used H3K4me3 data from an ABC cell line stored in the ENCODE database (GSE86718_ENCFF763KFL, replicated peaks BED file) for *peak integration* with significantly upregulated STAT3 peaks. Human genome version hg19 was used for analysis. RNA-seq data (GSE50721) from the same super-series was used for integration with STAT3 and H3K4me3 peaks, whereby we used statistical results provided by the authors of the study. For demonstrating the *nearby genes annotation* function, we used data from (56) and GEO dataset GSE52974, describing long-range enhancer elements regulating Myc expression for normal facial morphogenesis, mapped to mouse genome version mm9. For demonstration of the *TF & HM enrichment* function we used RNA-seq data from a longevity study focusing on daf-16/FoxO mutants in *C. elegans* ((57) and GEO dataset GSE72426) and *C. elegans* genome version ce11/WBcel245.

Analysed RNA-seq data from developing indirect flight muscle (IFM) in *Drosophila melanogaster* were taken from GSE107247 (46). We used the DESeq2 comparisons between time-points 72 h and 30 h after puparium formation (APF) provided by the authors. Genes with log2FC in absolute value greater than 2 and FDR smaller than 0.01 were considered as differentially expressed. For integration with phenotypic data, we made use of data published on muscle morphogenesis and function in *Drosophila* (58) and flight muscle growth in *Drosophila* (47).

For integrating gene expression changes of *yki* knocked-down flight muscles and flight muscles expressing a constitutively active *yki* compared to wild type, with Yki and Sd protein genome occupancy data, we used BRB-seq data from *yki* knock-down and *yki* constitutively active fly strains (47) available from GEO (GSE158957). Optimal IDR threshold narrow peak files for Yorkie (Yki, ENCSR422OTX) and Scalloped (Sd, ENCSR591PRH) ChIP-seq were downloaded from the ENCODE/modERN resource, both for *Drosophila* genome version dm6.

### *Drosophila* experimental methods

*Drosophila* strains were grown under standard conditions at 27°C to enhance GAL4 activity. All RNAi knock-downs were induced with *Mef2*-GAL4, a GAL4 line specifically expressed during development of all muscle types of the fly (58). UAS RNAi lines used were from the Vienna (59) and Harvard collections (60) and ordered from VDRC or Bloomington stock centres. See Supplementary Table S6 for all genotypes used. The flight test was done as described in (58).

Indirect flight muscle morphology was analysed in 3 to 5 days adult males or in 90 h APF pupae as previously published (61). Briefly, 90 h APF pupae or adult flies were fixed with 4% paraformaldehyde in PBT (PBS, 0.5% triton X-100) and cut into half-thoraces using a sharp microtome blade. The half-thoraces were blocked with 3% normal goat serum in PBT for 30 min and F-actin in the flight and leg muscles was visualised with phalloidin (coupled to Alexa488 or rhodamine, Molecular Probes, 1:1000 in PBT for 2 h or overnight) and Sls was stained with anti-Kettin antibody MAC155 (Babraham Institute, 1:100 in PBT overnight). The stained half-thoraces were imbedded in Vectashield (Biozol) and imaged on an LSM780 or LSM880 confocal microscope. Images were processed using Fiji (62).

## Results

### The AnnoMiner web server

AnnoMiner is composed of a *front-end*, for interacting with the user; and a *back-end,* which performs data upload, database maintenance and data analysis, as well as data visualization (Figure 1, Supplementary Figure S1).

**Figure 1:**
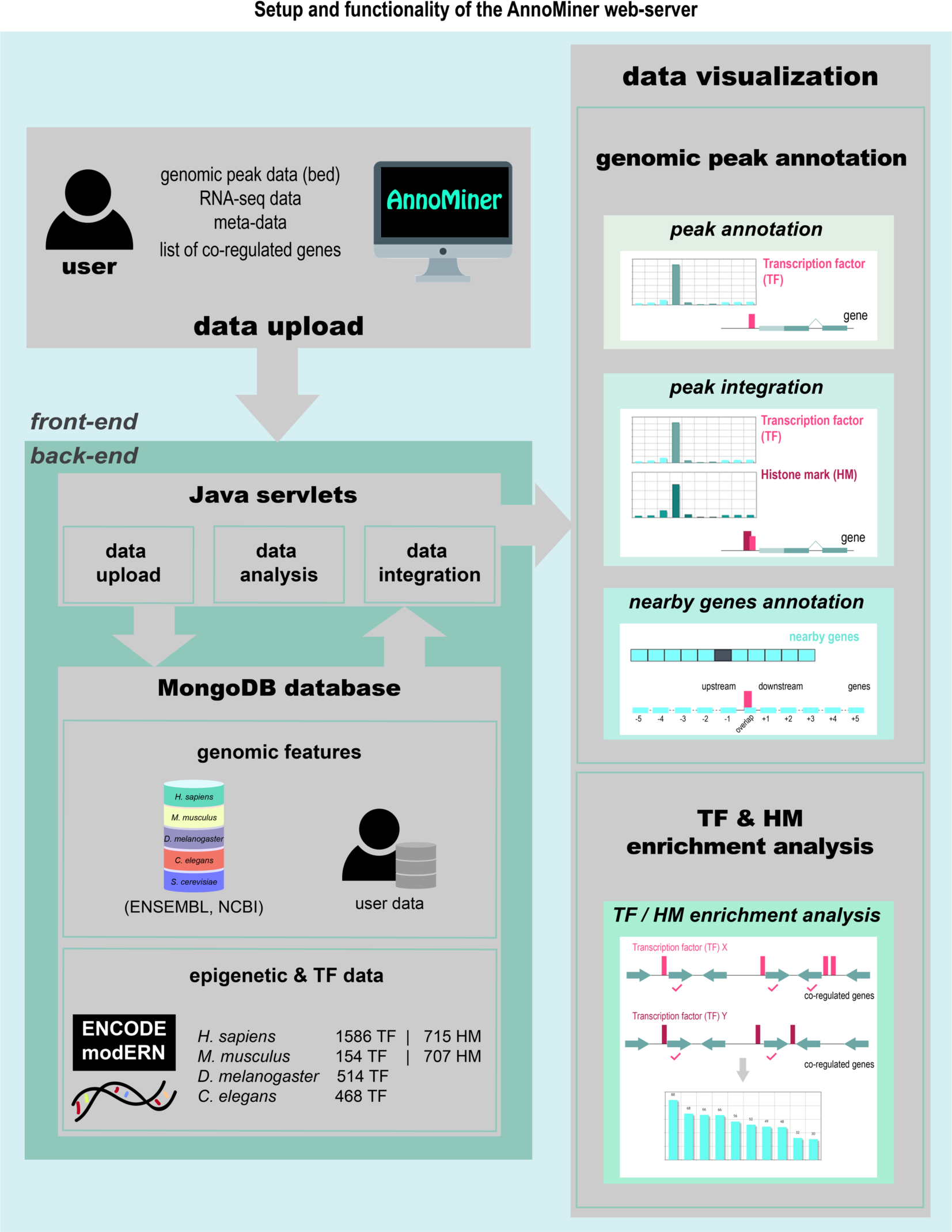
Setup of the AnnoMiner web-server. The user interacts with the web-server via the front-end for data upload and data mining. Genomic peak data (in BED format), annotation data, e.g. from transcriptomic analysis, or a list of co-regulated genes can be uploaded to the server. The user can either annotate peaks (*peak annotation*), integrate peaks (*peak integration*) or annotate nearby genes (*nearby genes annotation*) of a peak. AnnoMiner’s fourth function allows searching for enriched transcription factor binding or histone modification events in the promoter regions of a list of co-regulated genes (*TF & HM enrichment analysis*). In the back-end, Java servlets are responsible for data upload, analysis, integration and ID conversion. User data, genome feature data from ENSEMBL, UCSC and NCBI, as well as ENCODE, modENCODE and modERN ChIP-seq data are stored in a MongoDB database. Currently, we host genome feature data from *H. sapiens*, *M. musculus*, *D. melanogaster*, *C. elegans* and *S. cerevisiae*. Data from ENCODE, modENCODE and modERN are available in AnnoMiner for human, mouse, *Drosopihla* and *C. elegans*.

#### Web interface of AnnoMiner (front-end)

The web interface of AnnoMiner allows the user to upload, integrate and visualize the results. AnnoMiner’s web interface is intuitive and fully interactive relying on the JavaScript library JQuery and Bootstrap 4. Data upload, parameter settings and results are shown on the same page. To exploit the tool’s multiple functions, the front-end is sending JQuery’s POST requests to the back-end’s Java Servlets.

#### Back-end of AnnoMiner

The *back-end* of AnnoMiner is developed in Java and is running as a Java web server application deployed on Tomcat 8 on a Debian operating system. Separate Java Servlets were developed for each functionality, in order to be modular and easily implementable. The data used for the analysis are stored in a non-relational MongoDB database.

### Genomic peak annotation functions

Annominer’s three annotation functions (*peak annotation, peak integration and nearby genes annotation*, Figure 1) take as input one or more files containing genomic coordinates (in BED format), for example from ChIP-seq, with the aim of finding their associations with annotated gene features. This is done by determining the overlap between the genomic coordinates of a peak and the attributes of annotated gene features. The user can set the following parameters for the search: the minimum required overlap among the gene feature attributes and peak features (in bp or %); whether only the longest isoform of a gene or all its isoforms will be considered; the gene’s directionality; the organism and the genome resource. Optionally, a user-provided annotation file can be uploaded, containing for instance differential expression data which will be integrated with the gene lists generated from AnnoMiner annotation. The user-provided dataset is accepted in csv (comma separated values) or tsv (tab separated values) format. It can contain up to 6 columns, without any constraints in content, except the first column has to contain gene IDs to allow integration with AnnoMiner results.

#### Peak annotation

The *peak annotation* function computes the total coverage of the user-provided genomic regions (representing the peaks) with the attributes of each annotated gene feature in the genome assembly. AnnoMiner considers already annotated attributes of gene features (5’UTR, CDS and 3’UTR) as well as attributes or gene features provided by the user; in particular the promoter region is fully customizable with respect to the upstream and downstream region of the annotated TSS, as is the 5’ flanking region upstream and 3’ flanking regions downstream of the gene body (Figure 2 a). The first result shown by AnnoMiner is a coverage plot, visualizing the total base pair coverage of all peaks with the annotated attributes of the gene features (Figure 3 a). The user next chooses a target region (corresponding to a gene feature attribute) by clicking on one of the bars and on the ‘Show Genes!’ button (Figure 3 b). An interactive, sortable and downloadable table of all genes which overlap with the selected target region with peaks from the BED file is returned to the user (Figure 3 c). If the user provides a gene-based, custom annotation file, for instance containing differential expression data, these data will be integrated and displayed in the resulting table (Figure 3 d). While an annotation file will in most cases contain differential expression data based on RNA-seq analysis, it can contain any numerical or even text data. In summary, *peak annotation* in AnnoMiner annotates genomic regions provided by the user with gene features and their attributes in a flexible and user-centric way.

**Figure 2:**
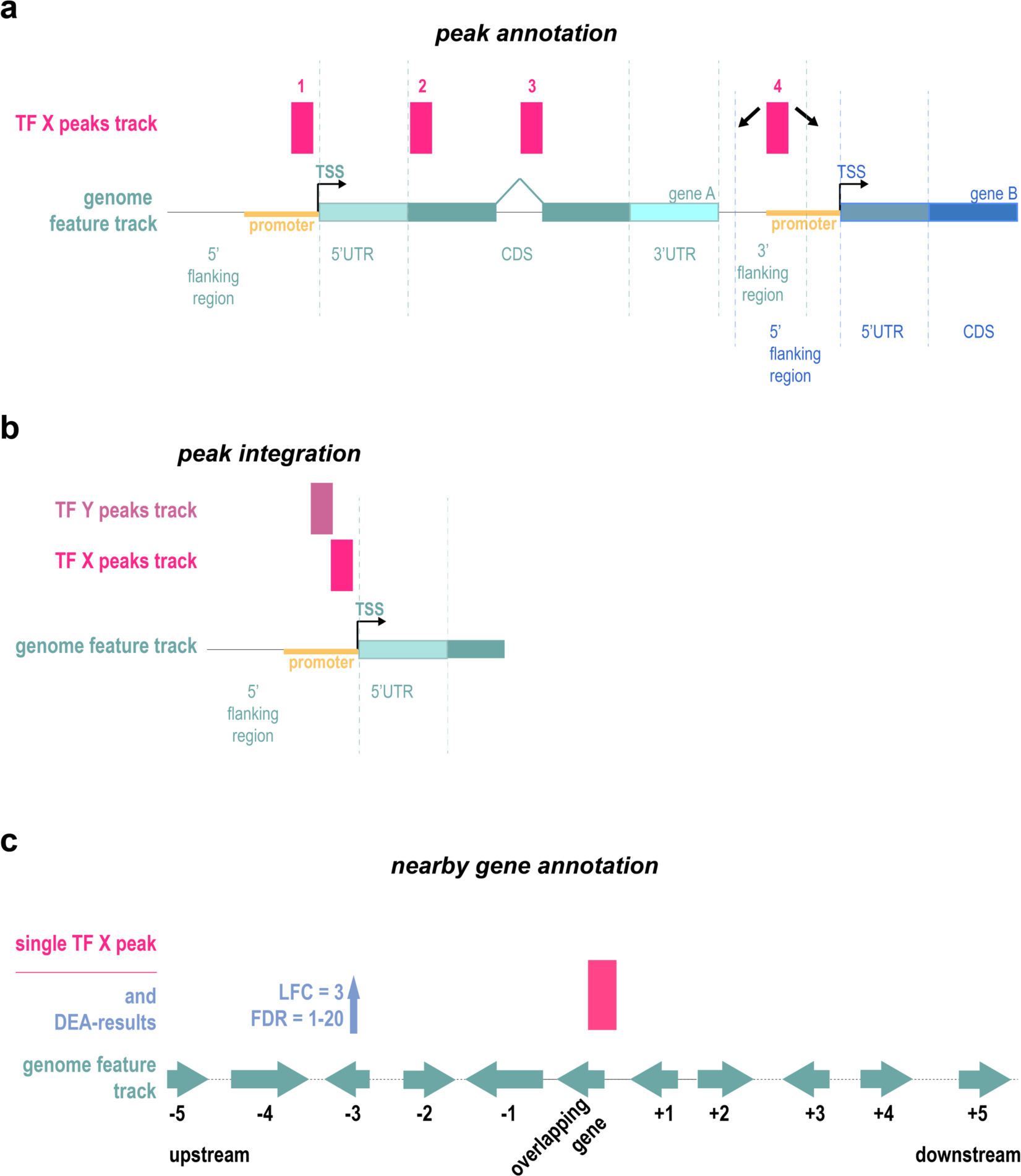
Different scenarios for the integration of peaks with gene features. **(a)** The peaks from a transcription factor (TF) track can be located in the promoter region or 5’ flanking region (peak 1) of gene A, in the coding region (peaks 2 and 3) or in the 3’ flanking region of gene A (peak 4). Peak 4 is at the same time located in the 5’ flanking region / promoter region of gene B. Its association is therefore not strictly clear, which should be considered during peak annotation. **(b)** The peaks of two transcriptional regulators (two TFs, two Histone Marks (HMs), or a TF and a HM, etc.) can be integrated based on their overlapping or nearby location in different attributes of a gene feature, as here demonstrated by their common overlap with the promoter region of a gene. **(c)** Activity of a transcriptional regulator, such as an enhancer, is not necessarily linked to the closest gene, but could affect neighbouring genes up- or downstream of the peak. Data from a differential expression analysis (DEA) of nearby genes can be correlated with a genomic peak harbouring a non-coding mutation or a genomic region harbouring a genome deletion.

**Figure 3:**
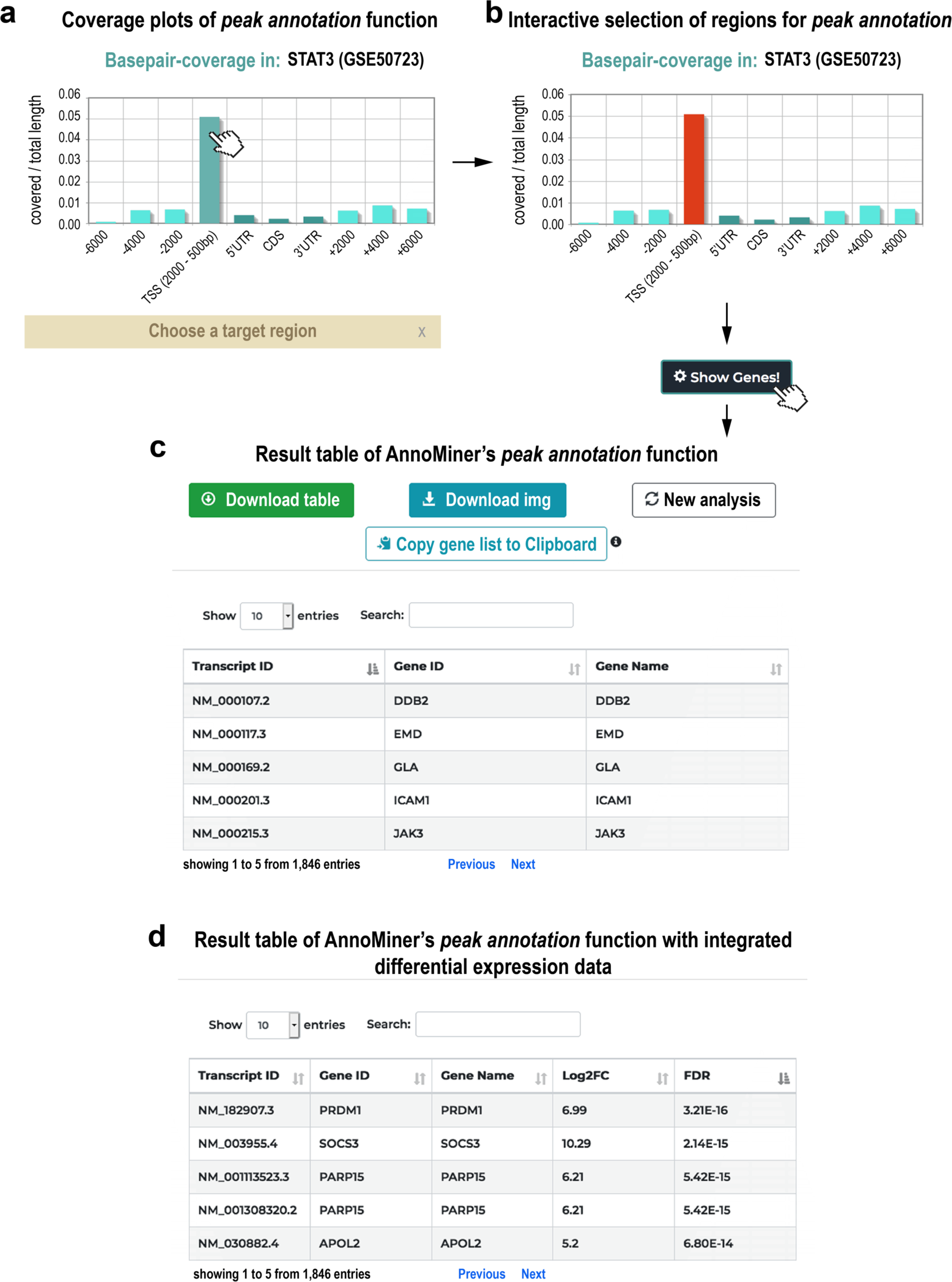
*Peak annotation* function of AnnoMiner. In the *peak annotation* function, AnnoMiner allows users to annotate peaks from a BED file. **(a)** After uploading a BED file to the server and choosing the function *peak annotation*, a coverage plot is produced, showing the covered base-pairs / total length relative to attributes extracted from the gene features of an organism’s genome. These include the 5’ upstream flanking region, the TSS (−2000bp and +500bp), the 5’UTR, coding sequence (CDS), 3’UTR, as well as the 3’ flanking region, whereby the boundaries for the TSS and flanking regions can be set by the user. **(b)** The plot is interactive and the user should select the target region (corresponding to the attribute of interest) to be analysed by selecting one or more bars to proceed. **(c)** When clicking on ‘Show Genes!’, the user retrieves a list of associated genes that fulfils the selected overlap criteria with the peaks. The table can be sorted according to different values, downloaded, browsed online, or the gene list can be copied to the clipboard for further analysis (e.g. enrichment analysis). The user can also select a new target region or start a completely new analysis. Significant differential peaks from STAT3 (GEO dataset GSE50723) were uploaded for producing this Figure. For **(d)**, differential expression data were uploaded from the associated GEO dataset GSE50721 from super-series GSE50724.

#### Peak integration

The *peak integration* function performs the *peak annotation* analysis, but for two genomic regions files (representing peaks from two independent TFs, a TF together with an HM or two independent HMs), allowing the user to integrate peaks from two different transcriptional regulators and identify gene features and their attributes that overlap with both (Figure 2 b). The same algorithm as in *peak annotation* is used for the annotation of each individual peak. With this function, genes co-regulated by two TFs can be identified, or a TF dataset can be integrated with chromatin structure data defined by histone marks, or other epigenetic information derived by other experimental techniques. Two coverage plots are returned by AnnoMiner (Figure 4 a), and the user chooses a target region for both and clicks on the ‘Show Genes!’ button (Figure 4 b). An interactive, sortable and downloadable table is returned with all genes that have a peak of both transcriptional regulators in their selected target regions (Figure 4 c). Custom gene annotation, such as differential expression data can again be provided, which is then integrated and displayed in the results table.

**Figure 4:**
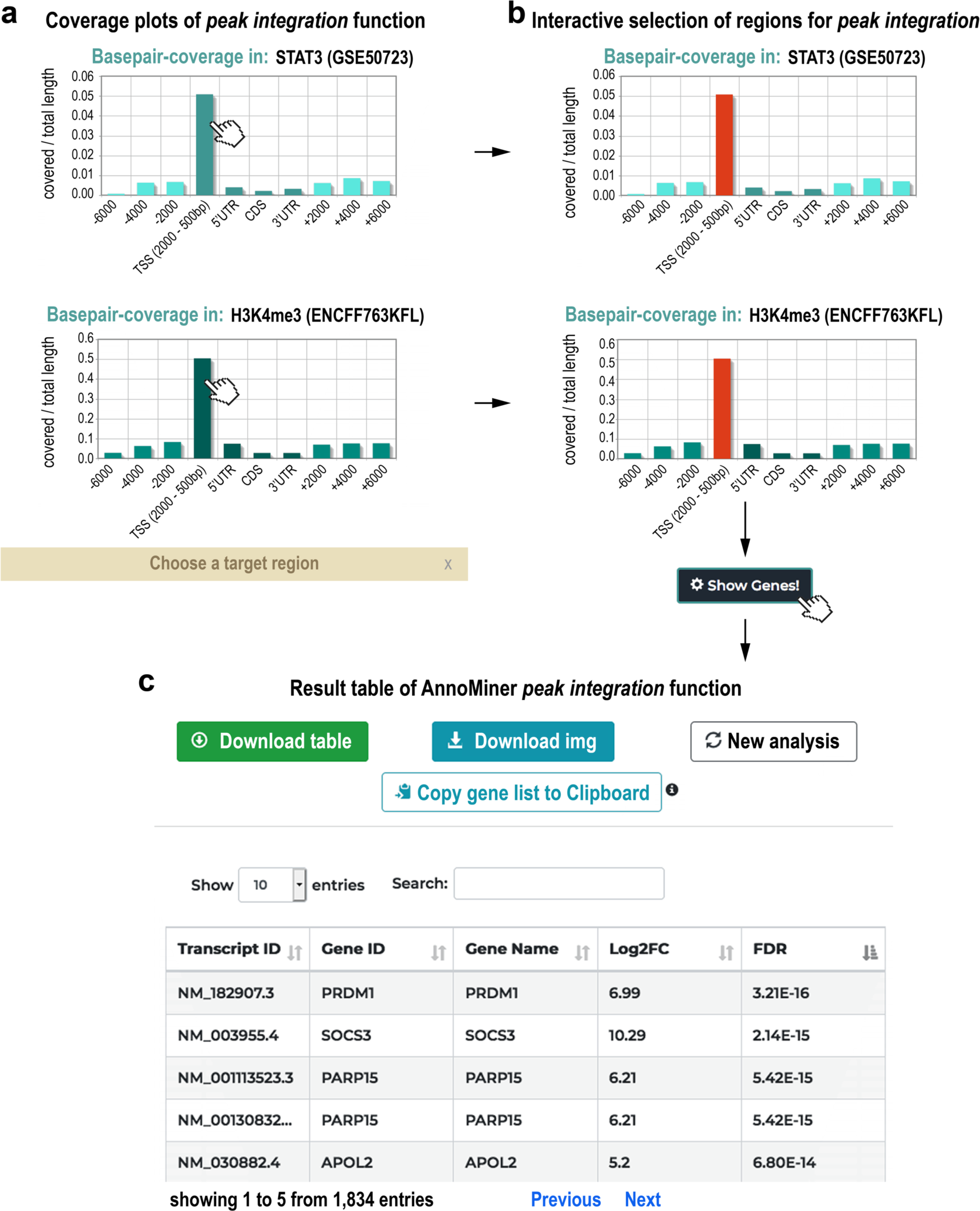
The *peak integration* function of AnnoMiner. If a user wants to integrate peaks from two transcriptional regulators and annotate them with gene feature attributes, the *peak integration* function must be chosen. **(a)** Two coverage plots of the two selected factors with gene feature attributes are shown to the user, from which a target region for each of them must be chosen **(b)**. **(c)** When clicking on ‘Show Genes!’, the associated genes are displayed as a table. Optionally, the user can upload a custom annotation file that contains for instance differential expression data (log2FC, FDR, etc.). These data are then shown in the resulting table. The table can be sorted according to different values, downloaded, browsed online, and the list of genes can be copied to the clipboard for further analysis. The user can also select a new target region or start a completely new analysis. Significantly upregulated peaks from STAT3 (GEO dataset GSE50723 from super-series GSE50724), as well as H3K4me3 methylation data from a comparable cell line (GEO dataset GSE86718) were uploaded for producing this Figure; differential expression data were taken from the associated GEO dataset GSE50721 from super-series GSE50724.

To show that AnnoMiner’s *peak integration* function can be used to integrate three different datasets in one analysis step, we re-analysed a data series published on STAT3 function in different forms of diffuse large B-cell lymphomas (DLBCL, GEO super-series GSE50724, (55)). Two subtypes of DLBCL are known, germinal centre B-cell-like (GCB) and activated B-cell-like (ABC). The ABC type responds only poorly to available therapies and can be often associated with an overexpression of STAT3 (55). The authors had compared STAT3 binding by ChIP-seq analysis between 8 patient-derived cell lines from GCB- and ABC type. They had performed RNA-seq analysis of the same cell lines to retrieve differentially expressed genes between the two subtypes. We made use of these data to identify genes with increased expression levels in the ABC type together with increased STAT3 binding events. We only considered STAT3 peaks that were significantly upregulated (FDR 0.05, fold change 1.25) in ABC-type DLBCL. To demonstrate the added value of Annominer’s *peak integration* function, we used H3K4me3 data from ENCODE from one of the ABC cell lines, OCI-Ly3 (accession: ENCFF763KFL), to limit the search to active promoters. Both peak files showed highest coverage with the direct promoter region of associated genes (Figure 4 a, b). We selected the major peaks for further analysis and integrated resulting peaks with differentially expressed genes from the same study. Of the upregulated unique genes, 42 contained an upregulated STAT3 peak in their promoter, as well as a H3K4me3 histone mark (Supplementary Figure S3 a, Supplementary Table S3). We submitted the list of 42 genes to the EnrichR web-server (44) and could identify terms strongly related to cancer, diffuse large B-cell lymphomas of the ABC type, IL10/STAT3 signalling and other relevant terms for the disease under study (Supplementary Figure S3 b, c; Supplementary Table S3). Moreover, AnnoMiner identified 28 additional direct targets of STAT3 to the ones already described in the original study (Supplementary Table S3). To summarize, AnnoMiner’s *peak integration* function was able to identify genes directly associated with ABC-type diffuse large B-cell lymphomas and potential direct targets for the transcription factor STAT3 by using a single user interface and a single AnnoMiner analysis step.

#### Nearby genes annotation

The *nearby genes annotation* function allows the user to visualize the differential regulation and retrieve the overlapping, as well as the closest 5 gene features up- and downstream of an individual peak (Figure 2 c). This function is most useful for exploring gene regulation in the vicinity of a genomic mutation or deletion in a non-coding region. Next to a BED file with the region of interest containing the mutation or genomic deletion, the user uploads an annotation file containing significantly differentially expressed genes. The resulting interactive AnnoMiner plot depicts the genomic neighbourhood of the peak, with the differential regulation of the overlapping and the five closest genes up- and downstream (Figure 5 a). The user can choose to visualize only the deregulation, or can discriminate between up- and down-regulation of the genes. In the latter case, the colour of the box reflects the direction of differential expression (either blue for up-, red for downregulated, green if an equal number of genes are up- and downregulated or grey if unchanged, Figure 5 a). By selecting one or more boxes (Figure 5 b), the selected genes, as well as their log2FC and FDR as uploaded by the user will be returned in a table (Figure 5 c).

**Figure 5:**
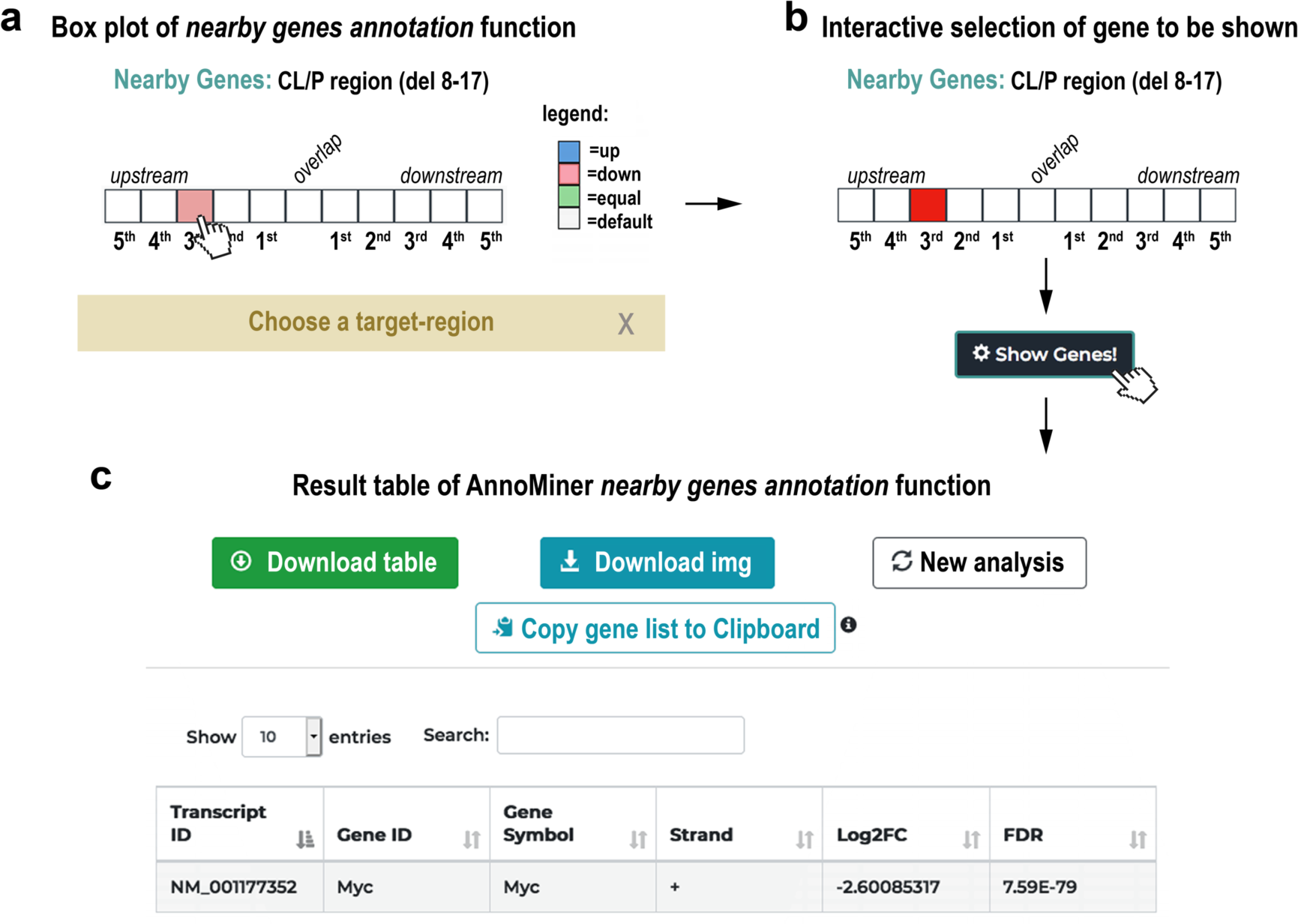
The *nearby genes annotation* function of AnnoMiner. **(a)** AnnoMiner’s *nearby gene annotation* function shows the differential regulation of 5 up- and down-stream, as well as the overlapping gene of a selected genomic peak. Next to a BED file with a single genomic region (a peak, a SNP or a genomic deletion), the user must upload an annotation file containing information on differential gene expression. Either general differential expression, or up- and down-regulation of neighbouring genes can be displayed. The user can also choose to consider directionality of the gene on the genomic strand. In the chosen example, a single gene is downregulated in the vicinity of the uploaded genomic regions file. **(b)** By selecting one or more boxes and pressing the ‘Show Genes!’ button, **(c)** the identity, as well as the log2FC and FDR of the selected gene(s) is returned in the table. We chose here data from a study that had shown the requirement of long-range enhancers regulating Myc expression for normal facial morphogenesis (GEO dataset GSE52974). Upon deletion of the medionasal enhancer (MNE) in mouse, Myc was the only gene observed to be differentially expressed in the vicinity of this deletion.

As a proof of concept for the *nearby genes annotation* function, we used a study that had shown the requirement of long-range enhancers regulating Myc expression for normal facial morphogenesis ((56), GEO dataset GSE52974). In humans, cleft lip or cleft palate (CL/P) is a frequent congenital malformation. This malformation has been associated with risk factors located at a 640kb noncoding region on chromosome 8. The corresponding region in mouse was studied by Uslu and colleagues and refined to a more specific enhancer region, the medionasal enhancer (MNE, 56). Deletions within the MNE in mouse led to smaller snouts and abnormalities of nasal and frontal bones amongst other defects. Myc was the only gene observed to be differentially expressed in the vicinity of this deletion. We used the CL/P deletion 8-17 (chr15:62668548 – 63550550) from Uslu et al. to create a single-peak BED file and uploaded it together with the significantly differentially expressed genes from the re-processed RNA-sequencing data from the same strain compared against control (GEO dataset GSE52974) to test the AnnoMiner *nearby genes annotation* function (Supplementary Table S4). Indeed, only a single gene is significantly differentially expressed (pink box, Figure 5 a), which is Myc (Figure 5 c). In principle, this function can also be used to explore the expression dynamics within the gene neighbourhood of multiple peaks (see Supplementary Figure S4). However, large-scale long-range gene regulatory data of this type are very sparse and their interpretation remains too complex to be exhaustively analysed with a tool like AnnoMiner. In summary, AnnoMiner’s *nearby genes annotation* function helps in an easy and quick way to identify deregulated genes in the neighbourhood of one genomic position.

#### Transcription factor & histone modification enrichment analysis

Transcription factor binding sites inferred from experimentally detected TF peaks in the genome can be used to predict TFs, which potentially co-regulate gene-sets. AnnoMiner’s *TF & HM enrichment analysis* function identifies enriched peaks in the promoter regions of a user-provided gene list, for instance co-regulated genes from a transcriptomic analysis. Any valid identifier is accepted, as AnnoMiner performs gene ID conversion on-the-fly using BioMart (51). AnnoMiner considers a gene as a potential target, if its promoter overlaps with a TF peak. The user can either choose the promoter region (up- and downstream number of base-pairs from the TSS) or use the *DynamicRanges* calculated by AnnoMiner, which is based on the distribution of a TF binding event relative to the TSS and therefore specific for each TF (see Methods, not available for histone marks). The results of the enrichment analysis are visualized as an interactive bar plot (for the first 10 hits, Figure 6 a), as well as an interactive table. In the table, all available TF ChIP-seq datasets in the AnnoMiner database for the species of interest are ranked according to their *Combined Score*. Along with this value, AnnoMiner also reports information about the experimental condition, cell line or developmental stage, contingency table values, p-value, enrichment score, FDR and the list of potential targets of the TF (Figure 6 b) in the downloadable version of the table.

**Figure 6:**
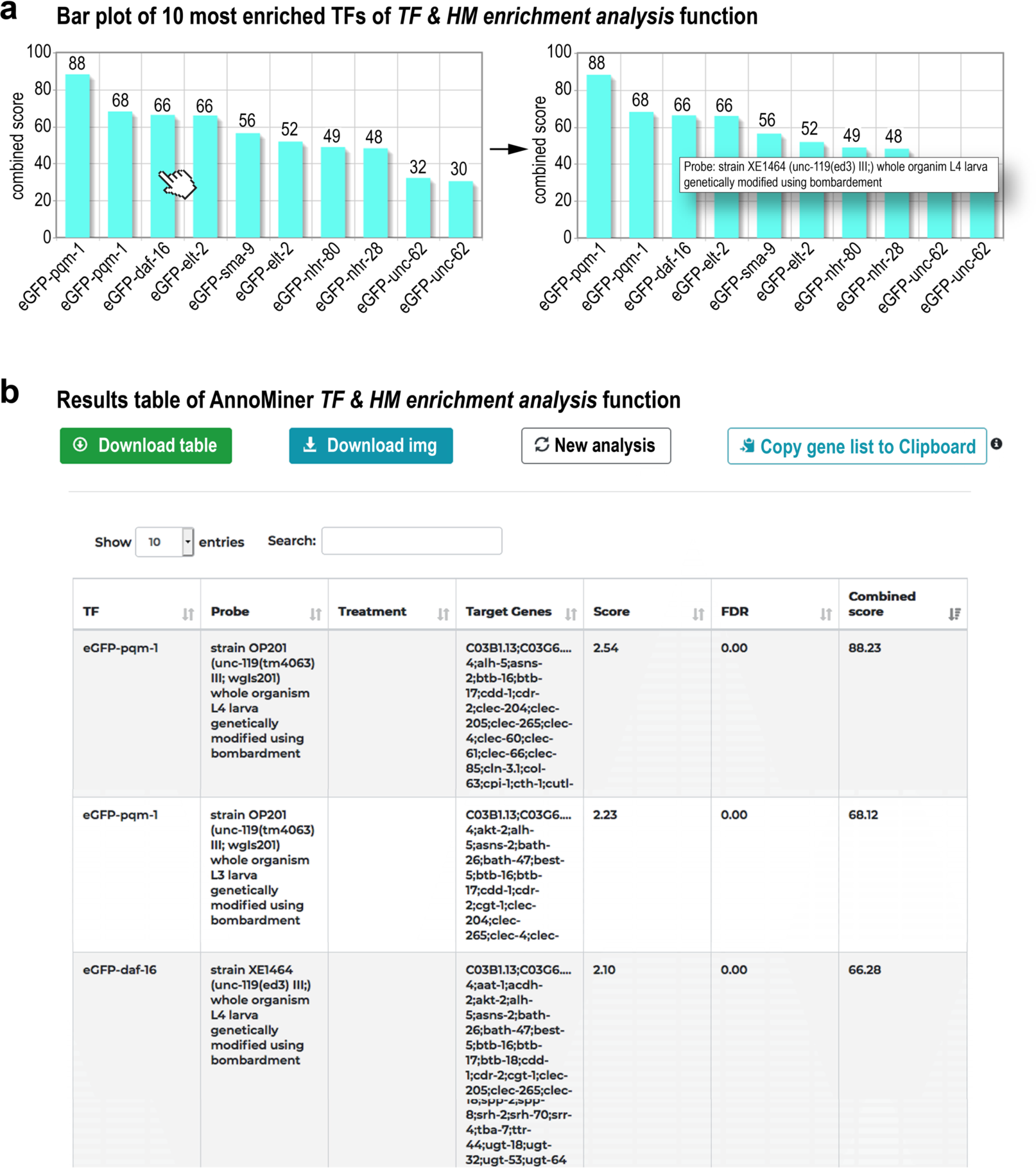
*TF & HM enrichment analysis* function of AnnoMiner. The output of a *TF & HM enrichment analysis* in AnnoMiner is first a plot of the top 10 enriched transcription factors **(a)**. The identity of the enriched TF with its combined score is shown. Hovering over the bar of a TF will reveal meta-information associated with the ChIP-seq dataset. **(b)** Second and at the same time, a table is returned, which contains the information on all TFs stored in the AnnoMiner database and their enrichments in the submitted gene list. The table is sortable and downloadable. Once downloaded, it includes information on the experimental design, treatment, the target gene for each TF, as well as the score, FDR and combined score. Target gene lists can be copied to the clipboard for further analysis. For demonstrating the *TF enrichment analysis* function of AnnoMiner, we took advantage of a study identifying direct daf-16/FoxO targets from a longevity study ((57) and GEO dataset GSE72426). In this study, the authors looked for isoform-specific targets of daf-16. We combined the list of all DAF-16 isoform targets and submitted it to AnnoMiner for identifying enriched transcription factors. AnnoMiner displays the 10 most enriched TFs, with their combined score indicated in the plot **(a)** as well as the table of all TFs stored for *C. elegans* and including enrichment information for the submitted gene list. Daf-16 is third hit; pqm-1 has been shown to bind to daf-16 response elements.

As a proof of principle for predicting transcriptional regulators we selected a differential expression dataset from *daf-16/FoxO* mutants in *C. elegans* (57). When uploading the list of all DAF-16A/F targets provided in (57) to AnnoMiner’s *TF & HM enrichment analysis* function, daf-16 was the 3^rd^ most significantly enriched transcription factor (Figure 6 a). Interestingly, pqm-1, which has been shown to bind to daf-16 response elements (63), was found at 1^st^ and 2^nd^ position by AnnoMiner. To conclude, AnnoMiner’s TF & HM enrichment function is a powerful tool for predicting relevant transcription factors co-regulating sets of genes with similar expression patterns.

#### Performance evaluation of AnnoMiner’s *TF & HM enrichment analysis* function

We wanted to compare the performance of AnnoMiner’s *TF & HM enrichment analysis* function with other web-tools for TF enrichment analysis. We followed in principle the evaluation protocol proposed by Keenan et al., which used PR-AUCs and ROC-AUCs calculated from the PPROC R-package for estimating performance ((43); for details see also Methods). In brief, we took manually curated datasets provided by (43) containing single TF perturbation experiments followed by RNA-seq from Gene Expression Omnibus (GEO (52)). Gene expression data used for benchmarking were restricted to experiments targeting TFs for which AnnoMiner is storing at least one high quality TF ChIP-seq dataset for the human assembly GRCh38, resulting in a total of 75 datasets that we could use for benchmarking. We submitted the list of significantly differentially expressed genes between perturbed TF versus wild-type control (which we hereafter refer to signature gene-sets) from these experiments to perform enrichment analysis using different tools, including AnnoMiner. We used the rank of the perturbed TF in the resulting enrichments of its associated signature gene-set to calculate PR-AUCs and ROC-AUCs. We furthermore calculated the cumulative distribution function for the ranks of each TF across all the experiments it was perturbed in. Only if a TF ranks randomly, the distribution function will be uniform; we performed Anderson-Darling tests to detect deviation from uniformity. We then computed the percentage of perturbed TFs that were correctly ranked within the first percentile to ensure that TFs were ranking high.

We chose the following web-tools for comparison: ChEA3, TFEA.ChIP and EnrichR (Table 1). ChEA3 offers the user 2 different methods to rank predicted TFs (meanRank and topRank) and we evaluated both ranking methods. EnrichR offers different resources for enrichment analysis and we used the resources ARCHS4, ChEA 2016, ENCODE 2015, ENCODE and ChEA Consensus and TRRUST 2019 for enrichment analysis, respectively. The number of datasets included in the benchmarkingSet depended on the resource tested (see Methods). The Anderson-Darling test returned significant results for all web-tools tested (ADtest in Table 1), except for EnrichR in combination with the ENCODE_and_ChEA_Consensus resource. This highlights the ability of all tools to rank the perturbed TF among the top candidates of the results. AnnoMiner outperformed TFEA.ChIP in all categories including percentage recovered TFs in the 1^st^ percentile (7.0 vs 0.0), ROC AU (0.69 vs 0.63) and PR AUC (0.68 vs 0.60). EnrichR differed in performance depending on the resource used. On average, it outperformed AnnoMiner on the percent recovered TFs (9.5 vs 7.0), while AnnoMiner reached slightly higher values in ROC AUCs (0.69 vs 0.66) and PR AUCs (0.69 vs 0.68). ChEA3 performed similar for both ranking methods used and outperformed AnnoMiner in all categories. To summarize, though it cannot reach the performance of ChEA, AnnoMiner outperforms the other evaluated tools in identifying relevant TFs at a high rank the in gene-sets derived from TF perturbation studies. Other than ChEA, however, AnnoMiner is available for all four major model organisms, including the invertebrates *Drosophila* and *C. elegans*.

### Using AnnoMiner to identify important transcriptional regulators of *Drosophila* flight muscle morphogenesis and growth

The assess the performance of AnnoMiner further, we tested its *TF & HM enrichment analysis* function to predict unknown transcriptional regulators for a list of co-regulated genes. We chose a dataset quantifying gene expression dynamic during development of *Drosophila melanogaster* indirect flight muscles (46), in which mRNA from indirect flight muscles had been isolated at several time-points correlating with key steps during muscle development (Figure 7 a). We focused on the development of the contractile apparatus called myofibrillogenesis and compared gene expression at 30 h after puparium formation (APF), when myofibrils assemble, with 72h APF, when myofibrils have matured (Figure 7 a). This comparison revealed 2193 differentially expressed genes that were shown in the original study to be strongly enriched for genes relevant for myofibril and mitochondrial development. We submitted this differential gene list to AnnoMiner (Supplementary Table S5) and searched all the 514 modERN and modENCODE TF ChIP-seq datasets from *Drosophila* stored in AnnoMiner for a potential enrichment of peaks. This identified 42 unique TFs significantly enriched with an FDR < 0.05. (Supplementary Table S5). The top 10 enriched datasets included Deaf1, Trl, Hr78 and cwo (Figure 7 b). Hence, these are potential transcriptional regulators of flight muscle development.

**Figure 7:**
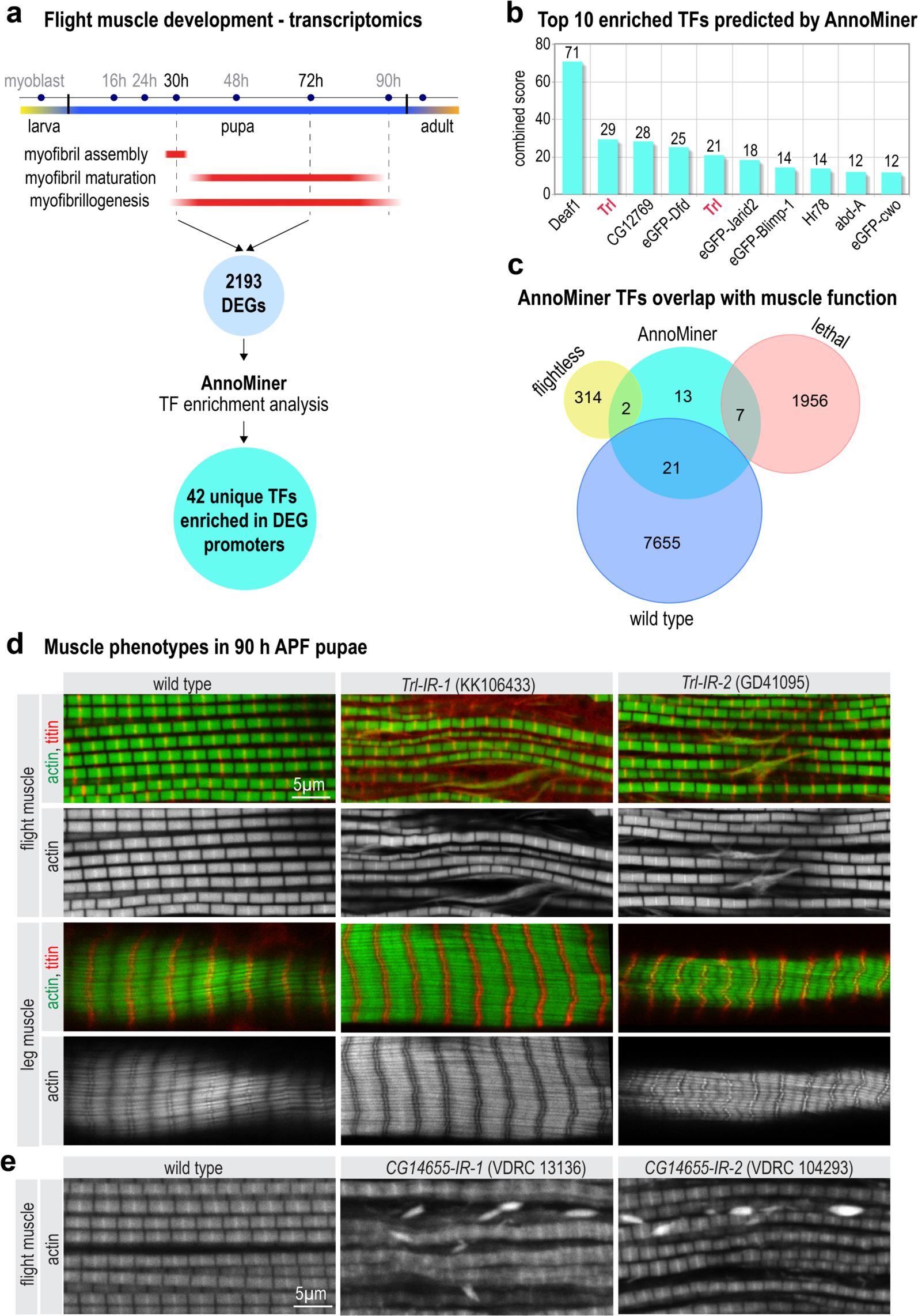
AnnoMiner identifies transcriptional regulators of flight muscle development. **(a)** Scheme of *Drosophila* indirect flight muscle development with different myofibrillogenesis phases highlighted in red. Comparing 30 h to 72 h APF revealed a list of 2193 differentially expressed genes (taken from GSE107247), which was uploaded to AnnoMiner. The *TF enrichment analysis* function of AnnoMiner identified 42 unique candidate TFs. **(b)** Output of the top-ten enriched TFs from AnnoMiner. Trl was identified from 2 different ChIP-seq datasets and is highlighted in red. **(c)** Phenotypic integration with muscle-specific RNAi knock-down data revealed candidate TFs with a potential function in muscle (flightless or lethal). Data on DEGs, phenotypic data from the RNAi screen, as well as annotated AnnoMiner enrichment results are available in Supplementary Tables S5. (**d)** *Trl* has a function during flight muscle myofibrillogenesis. Flight and leg muscles of 90 h APF pupae were fixed and stained for actin (phalloidin in green) and titin homolog Sls (anti-Kettin in red). Note that muscle-specific knock-down of *Trl* using independent hairpins (*Trl-IR-1* and *IR-2*) results in frayed flight muscle myofibrils compared to wild type control but rather normal leg muscle myofibrils. **(e)** *CG14655* has a function in flight muscles. Flight muscles of young adult flies from wild type or *CG14655-IR-1* or of 90 h APF pupae from *CG14655-IR-2* were stained for actin (phalloidin). Note the prominent actin accumulations present in both *CG14655* knock-down flight muscles.

To identify which of the 42 transcriptional regulators may have a function during flight muscle development we next integrated data from an RNAi-screen for muscle function (58), which had assayed for viability, flight muscle performance and body locomotion after muscle specific knock-down of individual TFs. From the 42 TFs identified by AnnoMiner, knock-down of two TFs resulted in flightless animals (*CG14655, cwo*), and seven were scored as lethal during development (*Trl, Hr78, lola, Vsx2, Pif1B, salr, Hr51*) (Supplementary Table S5); 21 TFs did not show a phenotype in this assay and the remaining 13 had not been tested (Figure 7 c).

#### Trithorax-like and the uncharacterized Zinc-finger protein CG14655 are required for flight muscle morphogenesis

For experimental verification we selected two proteins, Trl (Trithorax-like) and an uncharacterized zinc-finger protein called CG14655. We used muscle-specific knock-down to investigate a putative function of both genes in muscle. For *Trl* knock-down we used 4 independent transgenic RNAi lines driven with muscle-specific *Mef2*-GAL4. Two of those resulted in pupal lethality and the other two resulted in viable but flightless flies, demonstrating a function of *Trl* in flight muscle (Supplementary Table S6). For morphological analysis we visualized the myofibrils of flight and leg muscles of mature 90 h APF pupae in wild type and three different *Trl* knock-down lines. We found that knock-down of *Trl* caused disordered and frayed myofibrils in flight muscles, whereas leg muscle myofibrils appeared normal (Figure 7d, Supplementary Figure S5). This shows that Trl is required for normal myofibril development in flight muscle.

To investigate a role of *CG14655* during muscle development we also applied muscle-specific knock-down with *Mef2-*GAL4 and three different RNAi lines, one of which resulted in viable but flightless animals and two other overlapping hairpins resulted in pupal lethality (Supplementary Table S6). Morphological analysis showed that *CG14655* knock-down flight muscles displayed abnormal actin accumulations between their myofibrils, suggesting a role for *CG14655* in myofibril development of flight muscle. Together, these findings demonstrate the predictive power of AnnoMiner to identify transcriptional regulators by combining chromatin binding and differential expression data.

#### AnnoMiner helps identify targets co-regulated by Sd and Yki during flight muscle growth in Drosophila

Strikingly, two of the enriched transcriptional regulators in the above comparative flight muscle development dataset were the transcriptional effector of the Hippo pathway in *Drosophila* called Yorkie (Yki) and its essential Tead co-factor Scalloped (Sd) (64) (Supplementary Table S5). Recently, an essential function for the Hippo pathway promoting flight muscle growth by transcriptional up-regulation of mRNAs coding for sarcomeric proteins, which built the myofibrils, was identified (47). We wanted to know whether we could identify direct transcriptional targets of Yki/Sd during flight muscle growth. To this end, we integrated mRNA BRB-seq data from developing *yki* knock-down flight muscle (*yki-IR*), as well as from flight muscle expressing a constitutive active form of *yki* (*yki-CA)* compared to wild type controls (GEO accession GSE158957) with ChIP-seq data from Yki (modENCODE dataset ENCSR422OTX) and Sd (modERN dataset ENCSR591PRH) done in fly embryos using AnnoMiner’s *peak integration* function.

Both proteins showed prominent base pair coverage of the TSS regions of their target genes (Figure 8a). We selected these peaks to retrieve associated genes and then integrated the BRB-seq data for 24 h APF *yki* knockdown (*yki-IR* 24 h), as well as 24 h and 32 h APF constitutively active *yki*, respectively (*yki-CA* 24 h, *yki-CA* 32 h; Supplementary Table S7). Upon knock-down of *yki*, already at 32 h APF, a severe myofibril assembly defect had been observed (47). Interestingly, AnnoMiner identified Yki and Sd binding sites in the TSS of two genes essential for muscle function and development, which were downregulated in *yki* knock-down muscles at 24 h APF. This genes code for the sarcomeric proteins Tropomyosin 1 (Tm1) and the Nesprin-family protein Muscle-specific protein 300 kDa (Msp300), which is important to link the myofibrils to the nuclei (65) (Figure 8b). The gain-of-function *yki* phenotype (*yki*-*CA*) is characterized by premature expression of sarcomeric proteins resulting in muscle fiber hyper-compaction (47). At 24 h APF *scalloped* (*sd*) itself is the only direct target gene of the Yki/Sd complex which is differentially expressed in *yki-CA* (Figure 8c). At 32h AFP, AnnoMiner identified 177 unique genes and in total 541 transcripts as potential direct Yki/Sd targets. Using GO term enrichment analysis by modEnrichR (66) we found many processes and cellular compartments related to muscle development and function among the top enriched terms in these potential direct Yki/Sd targets (Figure 8 d, Supplementary Table S7). To conclude, using AnnoMiner’s *peak integration* function, we identified putative direct targets of the Yki/Sd transcriptional complex that showed differential expression upon *yki* knock-down or *yki* constitutive activation. Many of these genes are likely important for flight muscle morphogenesis.

**Figure 8:**
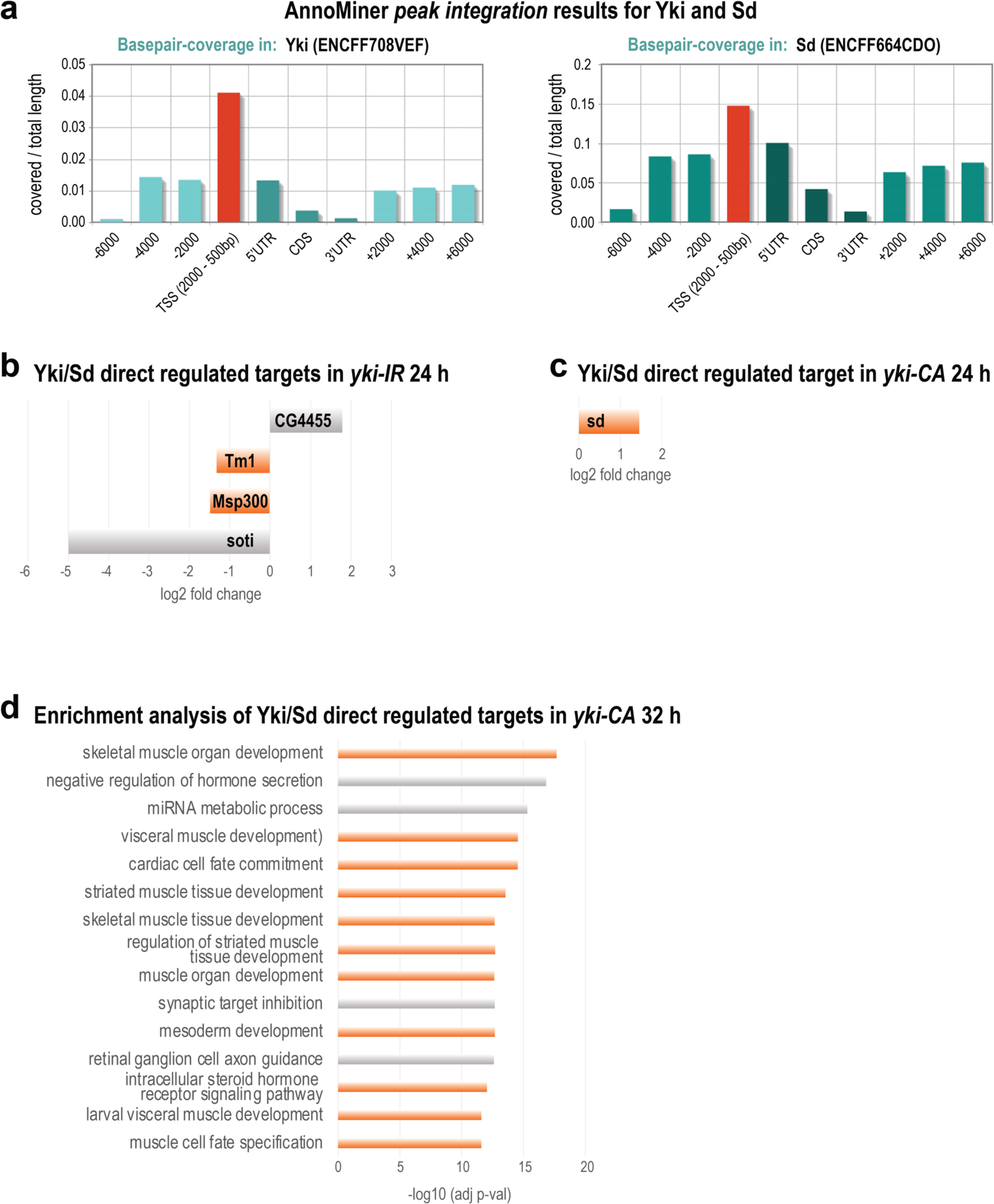
The Yorkie and Scalloped transcriptional complex regulates muscle-specific gene expression during flight muscle growth. **(a)** Using AnnoMiner’s *peak integration* function with ChIP-seq data from Yorkie (modENCODE ENCFF708VEF) and Scalloped (modERN ENCFF664CDO) we found high base-pair coverage of both TFs in the TSS (−2000bp - +500bp of the Transcription Start Site) of putative target genes. We selected these peaks from both TFs and integrated them with flight muscle specific BRB-seq data ((47) and GEO dataset GSE158957) isolated from a *yorkie* loss-of-function muscles at 24h APF **(b)** or yorkie gain-of-function muscles at 24h **(c)** and 32h APF **(d)**. The direct Yki/Sd targets *Tm1* and *Msp300* were downregulated in *yorkie* loss-of-function at 24 h APF. Only *scalloped* was identified as upregulated direct target from 24h *yorkie* gain of function muscles. At 32h, 177 genes were identified as potential direct Sd/Yki targets, many of which have muscle-related functions (see also Supplementary Table S6).

## Discussion

Here, we introduced AnnoMiner, a web-based, flexible and user-friendly platform for genomic peak annotation and integration, as well as transcription factor enrichment analysis. We illustrated AnnoMiner’s *peak annotation* and *integration*, as well as the *nearby genes annotation* functions with specific examples. We confirmed the predictive power of AnnoMiner’s TF enrichment function experimentally by identifying important regulators of *Drosophila* indirect flight muscle development. This was achieved searching for overrepresented TF peaks in promoters of genes differentially regulated during myofibrillogenesis using AnnoMiner.

AnnoMiner distinguishes itself from other peak annotation, as well as TF enrichment tools. AnnoMiner’s *peak annotation* and *peak integration* outputs first a bar plot that shows the overlap of a peak with different gene feature attributes, including up- and downstream regions, the TSS, 5’ and 3’ UTRs and the gene body. This has two advantages: first, the user can visualize the distribution of peaks of the uploaded file with respect to all relevant attributes of annotated gene features in the genome. Second, AnnoMiner allows to interactively choose the target region(s) for which the associated genes are returned. While other tools provide statistics on the peak distribution relative to gene feature attributes (e.g. (28)) in the output, to our knowledge, AnnoMiner is the only software that allows to easily retrieve specific gene-sets depending on the distribution of the peak coverage over gene feature attributes. The *peak integration* function offers the same flexibility. Moreover, both functions allow to directly integrate differential gene expression or other numerical data associated to genes with the genomic peak files. The *nearby genes annotation* function, which integrates expression data with peak data, is novel and for the first time, users can in a web-based manner visualize and retrieve genes that are not the nearest neighbours of a peak. It could be useful to integrate genomic, non-coding variants causative for human genetic diseases with disease-associated gene expression data. AnnoMiner’s *TF enrichment analysis* function offers to treat promoter regions dynamically for each specific TF with its *DynamicRanges* function. Finally, AnnoMiner is independent of the genomic assembly of the source data, as it on-the-fly translates submitted IDs and uses the ID compatible with the database chosen for gene centred peak annotation, as well as TF enrichment.

We compared AnnoMiner’s *TF enrichment analysis* function to the best-performing software in the field, which includes ChEA3, TFEA.ChIP and EnrichR. AnnoMiner could not reach the accuracy of ChEA3. One of the reasons could be that the dataset of TF - gene association used by ChEA3 supersedes data we retrieve from ENCODE, modENCODE and modERN, as it includes several additional datasets which are at least partially manually curated or generated. This hypothesis is supported by the fact that EnrichR shows differing performance when using different source data, showing lower performance when using the ENCODE data alone compared to the ones from ChEA3. One possible solution could be to add curated data to AnnoMiner’s *TF enrichment analysis* function, for instance from ReMap (67)(66). The disadvantage however is the higher cost in curation, as well as the fact that manually curated datasets are typically not available for model organisms such as *Drosophila* or *C. elegans*, but are rather restricted to human or mouse, as is the ChEA3 tool.

Finally, we predicted and verified potential transcriptional regulators of muscle and myofibril morphogenesis as well as muscle growth using AnnoMiner’s *TF & HM enrichment analysis* function. Amongst those is Trl, a GAGA transcription factor which contains a BTB/POZ domain, as well as a C2H2 zinc-finger that binds to DNA in a sequence-specific manner. Previous studies suggest that Trl is required to keep promoters nucleosome-free, thus allowing Pol-II-access (68). We showed here that muscle-specific knock-down of *Trl* using four independent hairpins either leads to pupal lethality or flightlessness. Consistently, we find severely perturbed indirect flight muscles upon *Trl* knock-down whereas leg muscles appear largely normal. This indicates a preferential function of Trl in flight muscle, however as 2 hairpins result in pupal lethality, a role of Trl in other body muscles is also likely.

A second potential direct transcriptional regulator identified by AnnoMiner is the uncharacterized Zinc-finger transcription factor CG14655. Muscle-specific knock-down of *CG14655* either results in pupal lethality or flightlessness and causes abnormal accumulations of actin in flight muscles. This again suggests that CG14655 is important for normal myofibril development in flight muscle.

Lastly, we made use of two other transcriptional regulators, Yorkie and Scalloped, which on DNA act in a complex (64), to identify its direct targets. AnnoMiner identified two direct targets of Yki, which upon loss of *yki* were downregulated. Both code for important muscle structure proteins, constituents of the sarcomere or linking the sarcomere to the nucleus, and hence could contribute to the severe phenotype observed upon *yki* knock-down. Gain-of-function of *yorkie* results in muscle fiber hyper-compaction and premature expression of sarcomeric protein components (47). Consistently, AnnoMiner identified a number of direct Yki targets with functions related to muscle development and growth. This substantiates a role for Scalloped and its transcriptional co-factor Yorkie during flight muscle growth.

## Conclusions

The new web-tool AnnoMiner is a user-friendly, intuitive, interactive and highly-flexible platform for genomic *peak annotation* and *peak integration*. It is suitable for identification of *nearby genes* of a genomic peak, as well as to perform *Transcription Factor and Histone Mark enrichment analysis* for a list of genes. This manuscript details all AnnoMiner functions and shows its usefulness for annotating and integrating peaks from two different ChIP-seq experiments together with transcriptomics data. Finally, AnnoMiner helped identify several key regulators of indirect flight muscle development and growth in *Drosophila*, some of which were confirmed experimentally.

## Acknowledgements

We thank Friedhelm Pfeiffer and all team members of the IBDM Computational Biology team for critical reading of the manuscript. We thank Aynur Kaya-Çopur for sharing data before publication. We acknowledge the France-BioImaging infrastructure supported by the French National Research Agency (ANR–10–INBS-04-01, Investments for the future). Fly stocks obtained from the Bloomington *Drosophila* Stock Center (NIH P40OD018537) and the Vienna *Drosophila* Resource Center (VDRC) were used in this study. This work was supported by the Turing Center of Living Systems (CENTURI, A*MIDEX, Investments for the Future) and the IBDM UMR 7288, by the Max Planck Society, the Centre National de la Recherche Scientifique (CNRS), and Aix-Marseille University.

## Availability of data and materials

The AnnoMiner web-server is freely available at http://chimborazo.ibdm.univ-mrs.fr/AnnoMiner/. The source code of AnnoMiner, as well as all data necessary to reproduce the study are available at https://gitlab.com/habermann_lab/annominer.

## Details on software provided in this manuscript

Project name: AnnoMiner

Project home page: http://chimborazo.ibdm.univ-mrs.fr/AnnoMiner/

Archived version: https://gitlab.com/habermann_lab/annominer

Operating system(s): Platform independent

Programming language: JavaScript, Java, PHP, MongoDB

Other requirements: none

License: GNU public license

Any restrictions to use by non-academics: none

## Author’s contributions

AM and FM were the main developers of the AnnoMiner web-server. FM performed all calculations, comparison with other tools and statistical analyses and together with BHH performed data analysis. MW and FS performed all experiments presented in this study. FM, FS and BHH wrote this manuscript, with contributions from AM. The project was conceived by BHH with contributions from AM and FM. All authors read and approved the final version of the manuscript.

## Ethics approval and consent to participate

Not applicable.

## Supplementary Data

Supplementary Figures S1-S5, Supplementary Tables S1, S2 and S6. (PDF format).

Supplementary Table S3 (Excel format).

Supplementary Table S4 (Excel format).

Supplementary Table S5 (Excel format).

Supplementary Table S7 (Excel format).

## Supplementary Data

### Supplementary Tables

**Supplementary Table S1:** Genome assemblies available in the AnnoMiner web-server.

| Assembly | Ensembl | Refseq | Ucsc | Genecode | Wormbase | Flybase |
| --- | --- | --- | --- | --- | --- | --- |
| hg38 |  | √ | √ | √ (34) |  |  |
| hg19 | √ | √ | √ | √ (34lift37) |  |  |
| mm10 |  | √ | √ | √ (M25) |  |  |
| mm9 | √ | √ (no mT) | √ |  |  |  |
| dm6 | √ | √ |  |  |  |  |
| dm3 | √ | √ (no mT) |  |  |  | √ |
| ce11 | √ | √ |  |  | √ (WS245) |  |
| sacCer3 | √ | √ |  |  |  |  |

**Supplementary Table S2:** Available TF ChIP-seq data for all model organisms and genome assemblies.

| Assembly | TF-ChIP-seq data available | unique TFs |
| --- | --- | --- |
| hg19 | 1687 | 666 |
| hg38 | 1586 | 660 |
| mm10 | 154 | 47 |
| dm6 | 514 | 474 |
| dm3 | 360 | 325 |
| ce11 | 468 | 286 |

**Supplementary Table S6:** Fly strains used and their observed phenotypes.

| Genotype | lethality | flight |
| --- | --- | --- |
| *Mef2-GAL4; UAS-GFP-Gma* / + | viable | wild type |
| *Mef2-GAL4; UAS-GFP-Gma* / *Trl-IR-1* (VDRC KK106433) | viable | flightless |
| *Mef2-GAL4; UAS-GFP-Gma* / *Trl-IR-2* (VDRC GD41095) | pupal lethal | - |
| *Mef2-GAL4; UAS-GFP-Gma* / *Trl-IR-3* (BDSC HMS02188) | viable | flightless |
| *Mef2-GAL4; UAS-GFP-Gma* / *Trl-IR-4* (VDRC GD17189) | pupal lethal | - |
| *Mef2-GAL4; UAS-GFP-Gma* / *CG14655-IR-1* (VDRC 13136) | viable | flightless |
| *Mef2-GAL4; UAS-GFP-Gma* / *CG14655-IR-2* (VDRC 104293) | pupal lethal | - |
| *Mef2-GAL4; UAS-GFP-Gma* / *CG14655-IR-3* (BDSC JF02334) | pupal lethal | - |

### Supplementary Figures

**Supplementary Figure S1:**
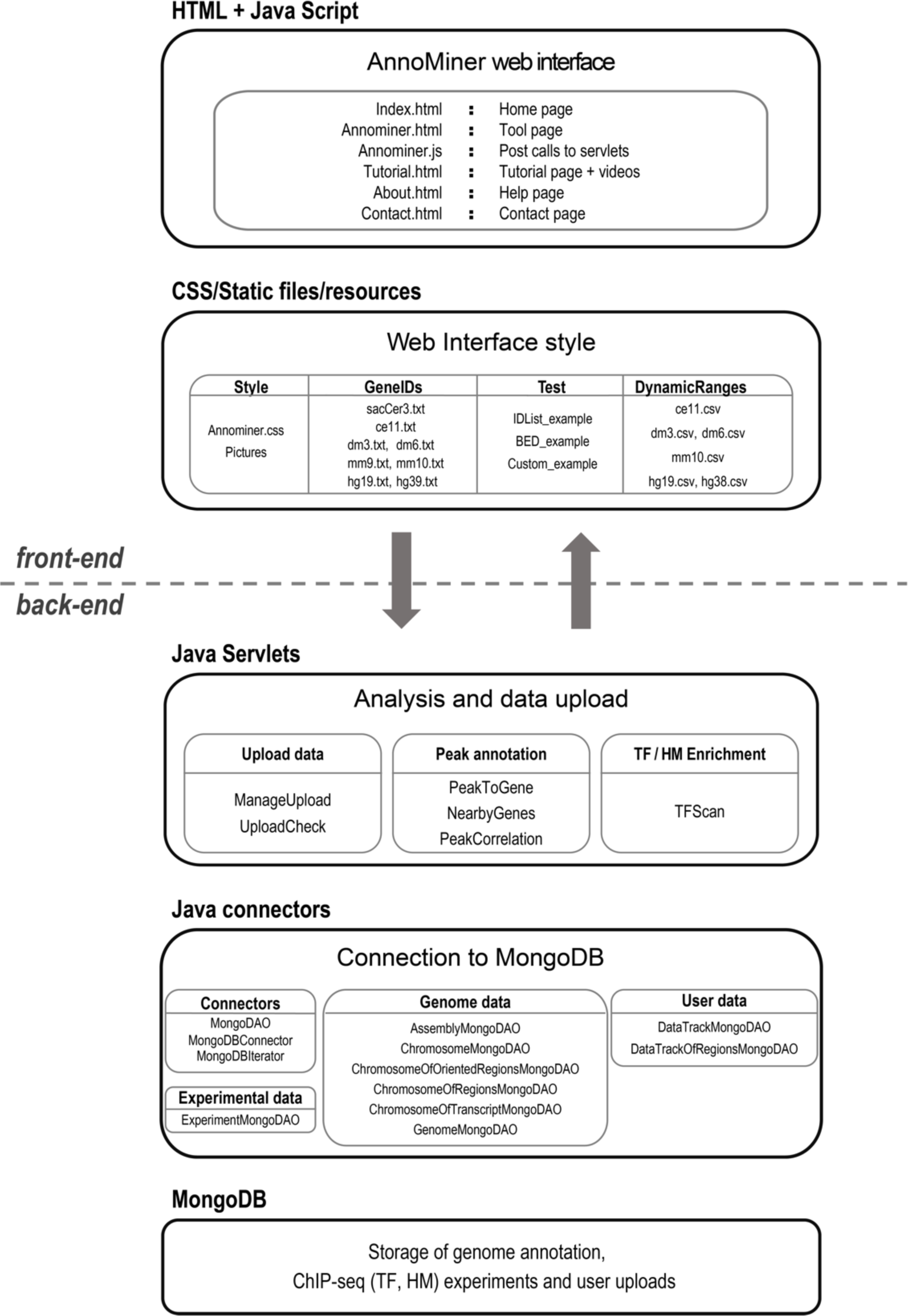
Back- and front-end structure of the AnnoMiner web-platform. In the back-end, the non-relational MongoDB database stores the annotated genomes, as well as the experimental epigenetic data (ChIP-seq data of transcription factors and histone modifications) that are publicly available via ENCODE, modENCODE and modERN. User-uploaded data are stored temporarily and are only available to the user. A set of java scripts connect to the MongoDB database for genome annotation and data retrieval. Different java servlets are created to carry out multiple functions in AnnoMiner: ManageUpload/UploadCheck to manage and store user data in the MongoDB; PeakToGene/NearbyGenes/PeakCorrelation to perform peak annotation and integration analysis; and TFScan to perform transcription factor (TF)/histone mark (HM) enrichment analysis functions. The front-end is made of static html pages where a main javascript file handles the DOM events, parameter choices and data visualizations. Among the static files are the GeneIDs files used for the gene id converter and the ones for the DynamicRanges which is needed by the TFScan function to dynamically assess the upstream promoter boundaries for each TF. Both these files are easily accessible and in plain text format in order to allow easy update and/or modification by the user. The interface appearance is set by the CSS layer (mainly Bootstrap 4). Finally, test files are provided, which together with the tutorial page and video tutorials aim to quickly get the user started in using AnnoMiner. To make AnnoMiner even more user-friendly, we have created tutorial videos, explaining in detail its usage, available at: http://chimborazo.ibdm.univ-mrs.fr/AnnoMiner/tutorial.html

**Supplementary Figure S2:**
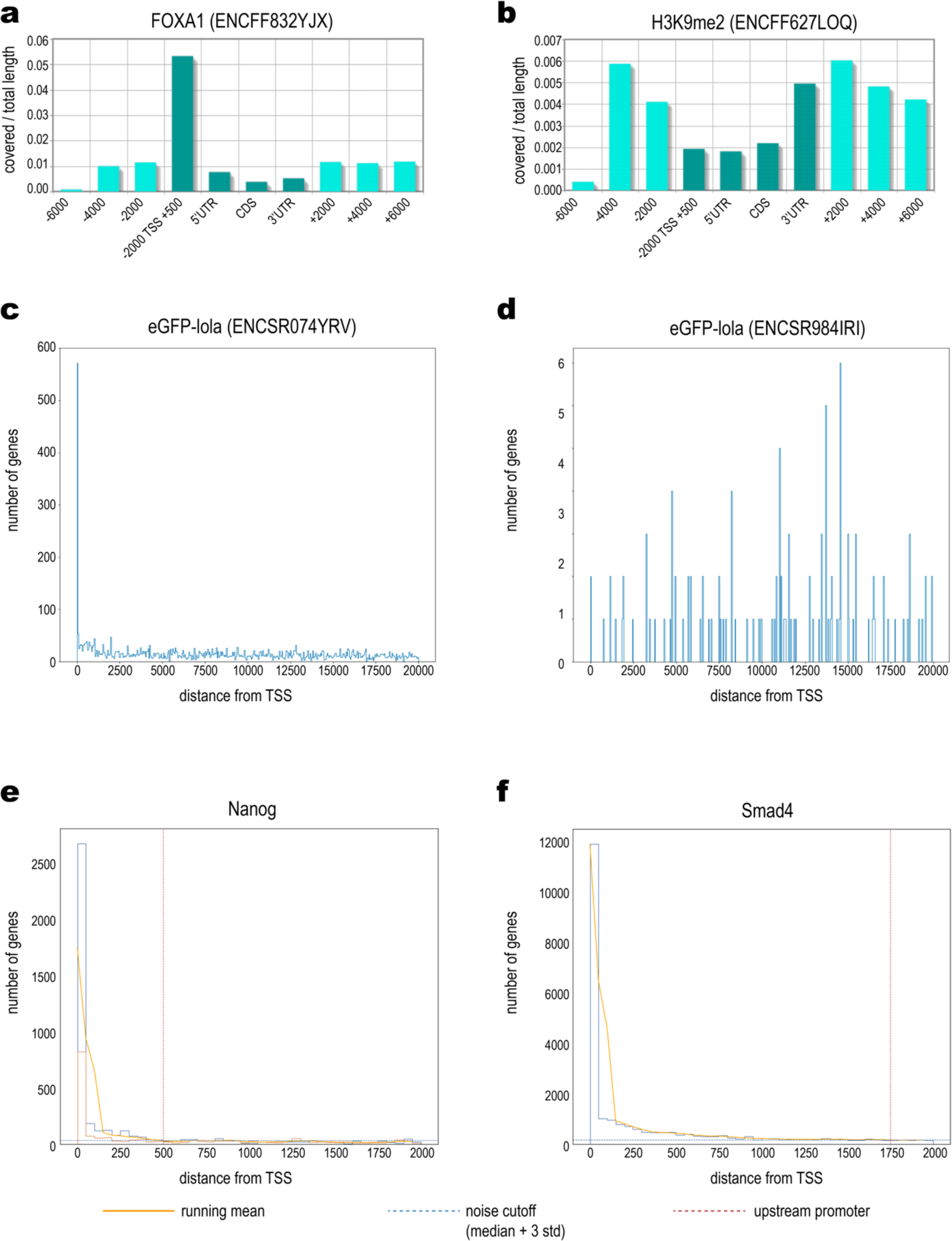
Coverage profiles of transcriptional regulators. **(a)** Coverage profile of the transcription factor FOXA1 (conservative IDR thresholded peaks downloaded from data set ENCSR735KEY, *H. sapiens* liver from a male adult, taken from the ENCODE database). A clear peak in the promoter region, defined as Transcription Start Site (TSS), flanked by −2000bp and +500bp respectively, can be seen. **(b)** Coverage profile of the transcriptional regulator TP53 (conservative IDR thresholded peaks taken from ENCODE data set ENCSR980EGJ, *H. sapiens* HepG2 cells). TP53 has a broader distribution, ranging from the TSS to −4000 bp upstream, as well as shows significant overlap with the 3’ downstream region of gene features. **(c)** Regular (ENCSR074YRV) and **(d)** outlier (ENCSR984IRI) distribution of the *Drosophila* transcriptional regulator Lola, available for dm6, Refseq. Distances between each peak and genes in a range of 20,000 bp upstream of their TSSs were computed, and then binned with a resolution of 50 bp. While ENCSR074YRV **(c)** clearly shows preferential binding close to genes TSS, ENCSR984IRI **(d)** shows few and non-preferential bindings. Therefore, it was marked and excluded as outlier. **(e)** Cumulative plot of the Nanog transcription factor over all uploaded data sets. Nanog’s running mean drops below the noise level at −500 bp upstream the TSSs of associated gene features. **(f)** Cumulative plot of the Smad4 transcription factor over all uploaded data sets. The running mean of Smad4 drops below the noise level only at −1750bp upstream of the TSSs of associated gene features. **(a + b)** Coverage is expressed as base pairs covered / total length.

**Supplementary Figure S3:**
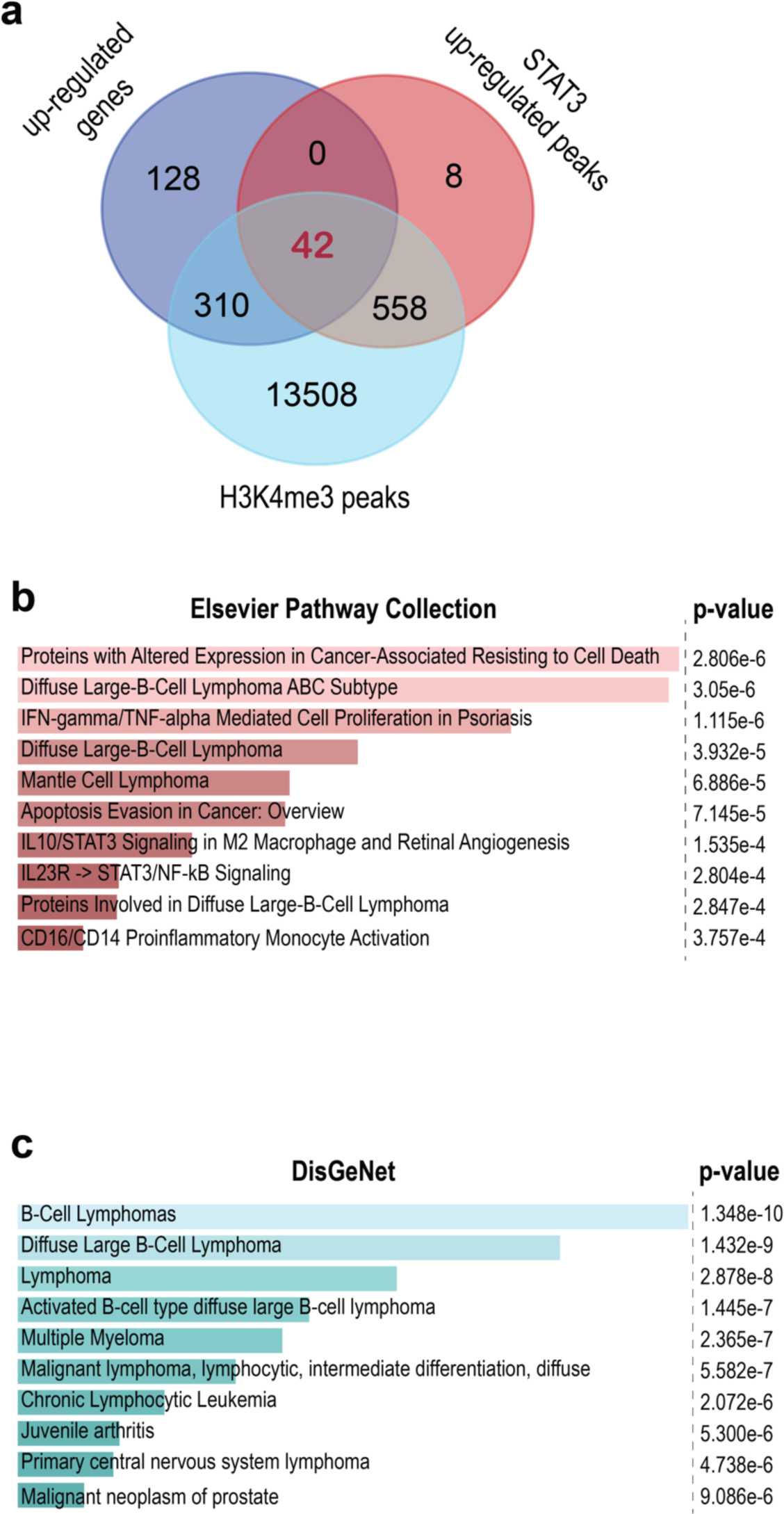
Data integration of STAT-3 upregulated peaks in different types diffuse large B-cell lymphomas (DLBCL) with H3K4me3 methylation status as well as differential expression data. TF data of STAT3 from two forms of DLBCL, the more aggressive activated B-cell like (ABC) with higher levels of STAT3 expression and the germinal center B-cell like (GBC) (GEO dataset GSE50723 from super-series GSE50724, 10.1534/g3.113.007674) were compared. Only peaks that were significant and upregulated in ABC with an FDR of <0.05 and a log2FC of >0.31 (corresponding to a fold change of >1.25) were chosen for upload. The major peak at the TSS shown in Figure 4 from the main text was chosen for further analysis. H3K4me3 methylation data from DLBCL cell line Ly3 was chosen to integrate STAT3 data with open chromatin status (GEO dataset GSE86718). Again, the significant peak at the TSS was chosen for further analysis (main Figure 4). **(a)** STAT3 upregulated peaks were integrated with H3K4me3 peaks, as well as upregulated genes in ABC subtype (taken from GEO dataset GSE50721 from super-series GSE50724). 42 genes were upregulated and had both, an upregulated STAT3 peak in ABC subtype and open chromatin in the cell line Ly3. These can be considered direct target genes of STAT3. **(b)** EnrichR enrichment analysis of these 42 intersecting genes revealed strong enrichment of pathway terms related to ‘Proteins with Altered Expression in Cancer-Associated Resisting to Cell Death’, ‘Diffuse Large-B-Cell Lymphoma ABC Subtype’, ‘Apoptosis Evasion in Cancer; Overview’ or ‘IL10/STAT3 Signaling in M2 Macrophage and Retinal Angiogenesis’, just to name a few. **(c)** DisGeNet enrichment using EnrichR revealed equally relevant terms, including ‘B-Cell Lymphomas’, ‘Diffuse Large-B-Cell Lymphoma’, Activated B-cell type diffuse large B-cell lymphoma’, among others. All data on upregulated genes associated with STAT3, H3K4me3 or both, as well as enrichment results are available in Supplementary Table S3.

**Supplementary Figure S4:**
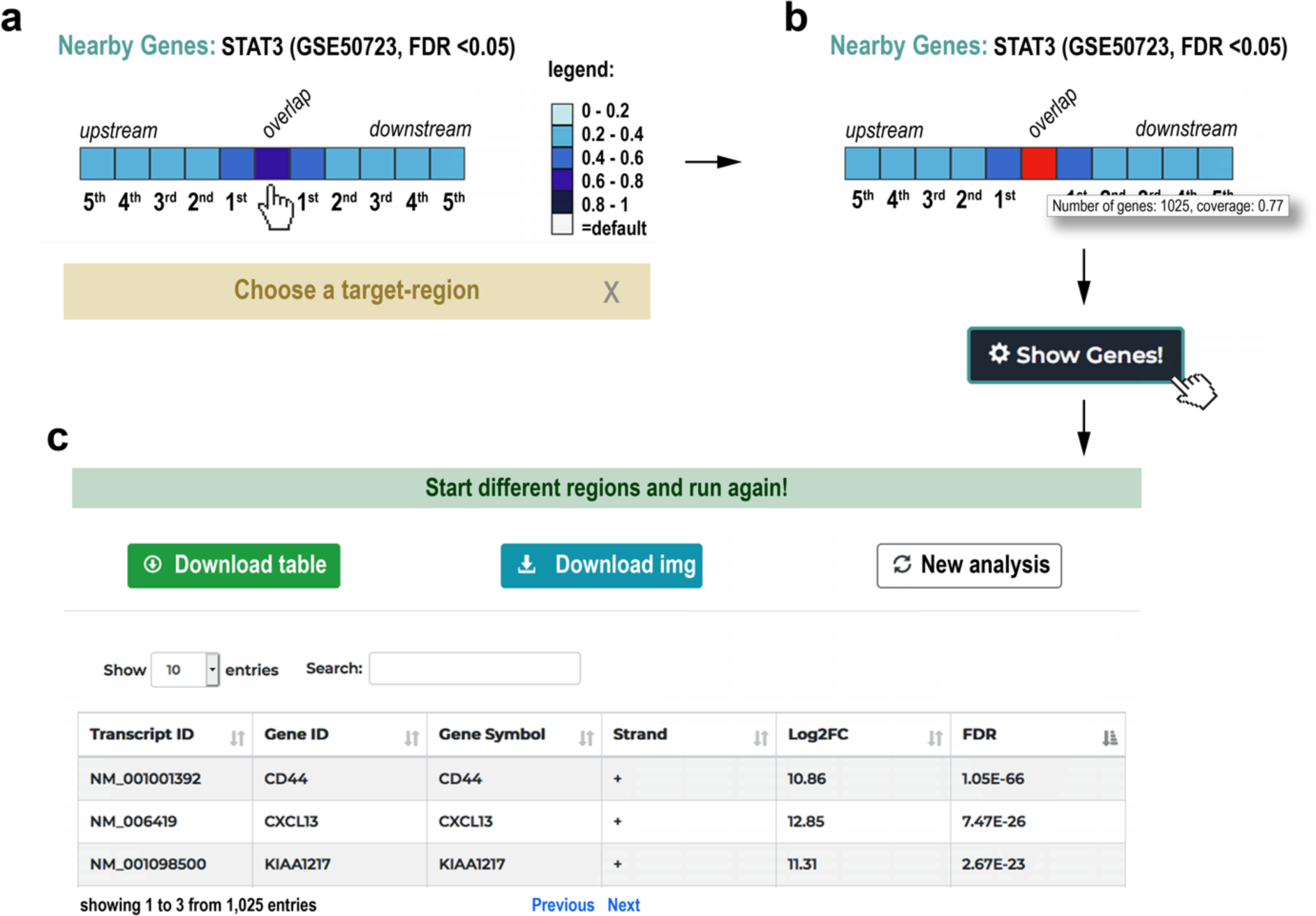
The nearby gene annotation function applied for multiple peaks. The nearby gene function can be also used to explore the correlation of multiple peaks of a TF with their 10 neighbouring genes. **(a)** The single-lane box plot shows the overlapping genes as well as the 5 upstream and 5 downstream neighbouring genes of all peaks from the submitted peak file. The colouring corresponds to the % deregulated genes at a certain position relative to the peaks. **(b)** The user can select a target region (here the overlapping genes), which pops up the information of associated genes and which percentage of those are deregulated. **(c)** When clicking on the ‘Show Genes!’ button, a table is shown with all deregulated genes of the selected region. Significant differential peaks from STAT3 (GEO dataset GSE50723) as well as associated differential expression data (GEO dataset GSE50721, both from super-series GSE50724) were uploaded for producing this Supplementary Figure.

**Supplementary Figure S5:**
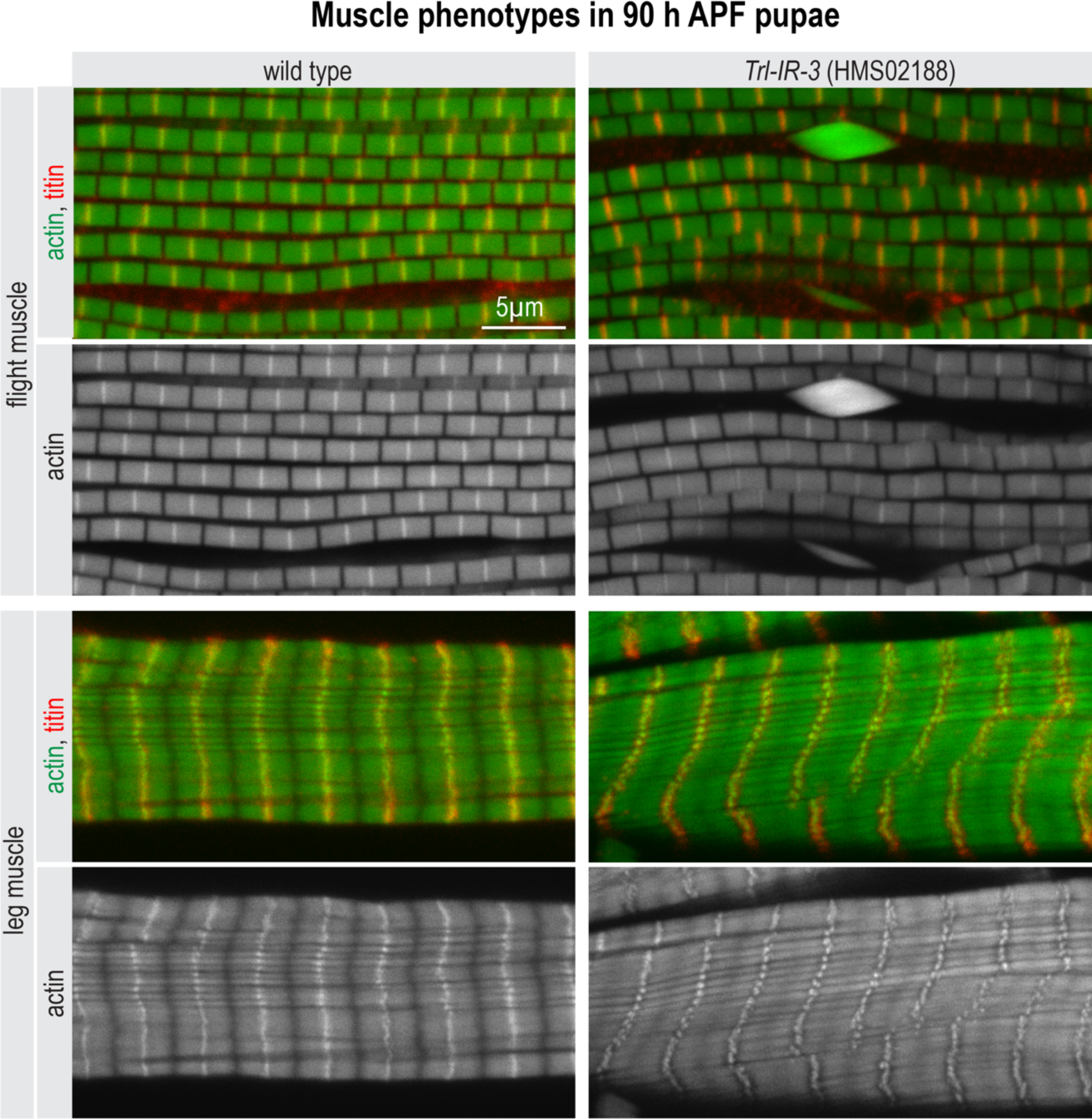
*Trl* has a function during flight muscle myofibrillogenesis. Flight and leg muscles of 90 h APF wild type or *Trl-IR-3* pupae were fixed and stained for actin (phalloidin in green) and titin homolog Sls (anti-Kettin in red). Note the abnormal actin accumulations in *Trl-IR-3* flight muscles compared to wild type.

## References

1. Zentner, G.E. and Henikoff, S. (2014) High-resolution digital profiling of the epigenome. Nat. Rev. Genet., 15, 814–827.

2. ENCODE Project Consortium, Snyder, M.P., Gingeras, T.R., Moore, J.E., Weng, Z., Gerstein, M.B., Ren, B., Hardison, R.C., Stamatoyannopoulos, J.A., Graveley, B.R., et al. (2020) Perspectives on ENCODE. Nature, 583, 693–698.

3. Schoenfelder, S. and Fraser, P. (2019) Long-range enhancer-promoter contacts in gene expression control. Nat. Rev. Genet., 20, 437–455.

4. Spitz, F. (2016) Gene regulation at a distance: From remote enhancers to 3D regulatory ensembles. Semin Cell Dev Biol, 57, 57–67.

5. Krivega, I. and Dean, A. (2012) Enhancer and promoter interactions-long distance calls. Curr. Opin. Genet. Dev., 22, 79–85.

6. Comoglio, F., Park, H.J., Schoenfelder, S., Barozzi, I., Bode, D., Fraser, P. and Green, A.R. (2018) Thrombopoietin signaling to chromatin elicits rapid and pervasive epigenome remodeling within poised chromatin architectures. Genome Res., 28, 295–309.

7. Mitchelmore, J., Grinberg, N.F., Wallace, C. and Spivakov, M. (2020) Functional effects of variation in transcription factor binding highlight long-range gene regulation by epromoters. Nucleic Acids Res., 48, 2866–2879.

8. Dao, L.T.M. and Spicuglia, S. (2018) Transcriptional regulation by promoters with enhancer function. Transcription, 9, 307–314.

9. Langmead, B. and Salzberg, S.L. (2012) Fast gapped-read alignment with Bowtie 2. Nat. Methods, 9, 357–359.

10. Nakato, R. and Shirahige, K. (2017) Recent advances in ChIP-seq analysis: from quality management to whole-genome annotation. Brief. Bioinformatics, 18, 279–290.

11. Furey, T.S. (2012) ChIP-seq and beyond: new and improved methodologies to detect and characterize protein-DNA interactions. Nat. Rev. Genet., 13, 840–852.

12. Thomas, R., Thomas, S., Holloway, A.K. and Pollard, K.S. (2017) Features that define the best ChIP-seq peak calling algorithms. Brief. Bioinformatics, 18, 441–450.

13. Wilbanks, E.G. and Facciotti, M.T. (2010) Evaluation of algorithm performance in ChIP-seq peak detection. PLoS ONE, 5, e11471.

14. Zhang, Y., Liu, T., Meyer, C.A., Eeckhoute, J., Johnson, D.S., Bernstein, B.E., Nusbaum, C., Myers, R.M., Brown, M., Li, W., et al. (2008) Model-based analysis of ChIP-Seq (MACS). Genome Biol, 9, R137–9.

15. Xu, H., Wei, C.-L., Lin, F. and Sung, W.-K. (2008) An HMM approach to genome-wide identification of differential histone modification sites from ChIP-seq data. Bioinformatics, 24, 2344–2349.

16. Huang, W., Umbach, D.M., Vincent Jordan, N., Abell, A.N., Johnson, G.L. and Li, L. (2011) Efficiently identifying genome-wide changes with next-generation sequencing data. Nucleic Acids Res., 39, e130–e130.

17. Shen, L., Shao, N.-Y., Liu, X., Maze, I., Feng, J. and Nestler, E.J. (2013) diffReps: detecting differential chromatin modification sites from ChIP-seq data with biological replicates. PLoS ONE, 8, e65598.

18. Ambrosini, G., Dreos, R., Kumar, S. and Bucher, P. (2016) The ChIP-Seq tools and web server: a resource for analyzing ChIP-seq and other types of genomic data. BMC Genomics, 17, 938–15.

19. Blahnik, K.R., Dou, L., O’Geen, H., McPhillips, T., Xu, X., Cao, A.R., Iyengar, S., Nicolet, C.M., Ludäscher, B., Korf, I., et al. (2010) Sole-Search: an integrated analysis program for peak detection and functional annotation using ChIP-seq data. Nucleic Acids Res., 38, e13–e13.

20. Guzman, C. and D’Orso, I. (2017) CIPHER: a flexible and extensive workflow platform for integrative next-generation sequencing data analysis and genomic regulatory element prediction. BMC Bioinformatics, 18, 363–16.

21. Boeva, V., Lermine, A., Barette, C., Guillouf, C. and Barillot, E. (2012) Nebula--a web-server for advanced ChIP-seq data analysis. Bioinformatics, 28, 2517–2519.

22. Salmon-Divon, M., Dvinge, H., Tammoja, K. and Bertone, P. (2010) PeakAnalyzer: genome-wide annotation of chromatin binding and modification loci. BMC Bioinformatics, 11, 415–12.

23. Quinlan, A.R. and Hall, I.M. (2010) BEDTools: a flexible suite of utilities for comparing genomic features. Bioinformatics, 26, 841–842.

24. Heinz, S., Benner, C., Spann, N., Bertolino, E., Lin, Y.C., Laslo, P., Cheng, J.X., Murre, C., Singh, H. and Glass, C.K. (2010) Simple combinations of lineage-determining transcription factors prime cis-regulatory elements required for macrophage and B cell identities. Mol. Cell, 38, 576–589.

25. Yu, G., Wang, L.-G. and He, Q.-Y. (2015) ChIPseeker: an R/Bioconductor package for ChIP peak annotation, comparison and visualization. Bioinformatics, 31, 2382–2383.

26. Kondili, M., Fust, A., Preussner, J., Kuenne, C., Braun, T. and Looso, M. (2017) UROPA: a tool for Universal RObust Peak Annotation. Sci Rep, 7, 2593–12.

27. Tang, X., Srivastava, A., Liu, H., Machiraju, R., Huang, K. and Leone, G. (2017) annoPeak: a web application to annotate and visualize peaks from ChIP-seq/ChIP-exo-seq. Bioinformatics, 33, 1570–1571.

28. Chen, T.-W., Li, H.-P., Lee, C.-C., Gan, R.-C., Huang, P.-J., Wu, T.H., Lee, C.-Y., Chang, Y.-F. and Tang, P. (2014) ChIPseek, a web-based analysis tool for ChIP data. BMC Genomics, 15, 539–13.

29. Huang, W., Loganantharaj, R., Schroeder, B., Fargo, D. and Li, L. (2013) PAVIS: a tool for Peak Annotation and Visualization. Bioinformatics, 29, 3097–3099.

30. Bhasin, J.M. and Ting, A.H. (2016) Goldmine integrates information placing genomic ranges into meaningful biological contexts. Nucleic Acids Res., 44, 5550–5556.

31. McLean, C.Y., Bristor, D., Hiller, M., Clarke, S.L., Schaar, B.T., Lowe, C.B., Wenger, A.M. and Bejerano, G. (2010) GREAT improves functional interpretation of cis-regulatory regions. Nat. Biotechnol., 28, 495–501.

32. Zhu, L.J., Gazin, C., Lawson, N.D., Pagès, H., Lin, S.M., Lapointe, D.S. and Green, M.R. (2010) ChIPpeakAnno: a Bioconductor package to annotate ChIP-seq and ChIP-chip data. BMC Bioinformatics, 11, 237–10.

33. ENCODE Project Consortium (2012) An integrated encyclopedia of DNA elements in the human genome. Nature, 489, 57–74.

34. Davis, C.A., Hitz, B.C., Sloan, C.A., Chan, E.T., Davidson, J.M., Gabdank, I., Hilton, J.A., Jain, K., Baymuradov, U.K., Narayanan, A.K., et al. (2018) The Encyclopedia of DNA elements (ENCODE): data portal update. Nucleic Acids Res., 46, D794–D801.

35. Jou, J., Gabdank, I., Luo, Y., Lin, K., Sud, P., Myers, Z., Hilton, J.A., Kagda, M.S., Lam, B., O’Neill, E., et al. (2019) The ENCODE Portal as an Epigenomics Resource. Curr Protoc Bioinformatics, 68, e89.

36. Gerstein, M.B., Lu, Z.J., Van Nostrand, E.L., Cheng, C., Arshinoff, B.I., Liu, T., Yip, K.Y., Robilotto, R., Rechtsteiner, A., Ikegami, K., et al. (2010) Integrative analysis of the Caenorhabditis elegans genome by the modENCODE project. Science, 330, 1775–1787.

37. Kudron, M.M., Victorsen, A., Gevirtzman, L., Hillier, L.W., Fisher, W.W., Vafeados, D., Kirkey, M., Hammonds, A.S., Gersch, J., Ammouri, H., et al. (2018) The ModERN Resource: Genome-Wide Binding Profiles for Hundreds of Drosophila and Caenorhabditis elegans Transcription Factors. Genetics, 208, 937–949.

38. Alvarez, M.J., Shen, Y., Giorgi, F.M., Lachmann, A., Ding, B.B., Ye, B.H. and Califano, A. (2016) Functional characterization of somatic mutations in cancer using network-based inference of protein activity. Nat. Genet., 48, 838–847.

39. Garcia-Alonso, L., Iorio, F., Matchan, A., Fonseca, N., Jaaks, P., Peat, G., Pignatelli, M., Falcone, F., Benes, C.H., Dunham, I., et al. (2018) Transcription Factor Activities Enhance Markers of Drug Sensitivity in Cancer. Cancer Res, 78, 769–780.

40. Wang, Z., Civelek, M., Miller, C.L., Sheffield, N.C., Guertin, M.J. and Zang, C. (2018) BART: a transcription factor prediction tool with query gene sets or epigenomic profiles. Bioinformatics, 34, 2867-2869.

41. Kwon, A.T., Arenillas, D.J., Worsley Hunt, R. and Wasserman, W.W. (2012) oPOSSUM-3: advanced analysis of regulatory motif over-representation across genes or ChIP-Seq datasets. G3 (Bethesda), 2, 987-1002.

42. Puente-Santamaria, L., Wasserman, W.W. and Del Peso, L. (2019) TFEA.ChIP: a tool kit for transcription factor binding site enrichment analysis capitalizing on ChIP-seq datasets. Bioinformatics, 35, 5339–5340.

43. Keenan, A.B., Torre, D., Lachmann, A., Leong, A.K., Wojciechowicz, M.L., Utti, V., Jagodnik, K.M., Kropiwnicki, E., Wang, Z. and Ma’ayan, A. (2019) ChEA3: transcription factor enrichment analysis by orthogonal omics integration. Nucleic Acids Res., 47, W212–W224.

44. Kuleshov, M.V., Jones, M.R., Rouillard, A.D., Fernandez, N.F., Duan, Q., Wang, Z., Koplev, S., Jenkins, S.L., Jagodnik, K.M., Lachmann, A., et al. (2016) Enrichr: a comprehensive gene set enrichment analysis web server 2016 update. Nucleic Acids Res., 44, W90–7.

45. Imrichová, H., Hulselmans, G., Atak, Z.K., Potier, D. and Aerts, S. (2015) i-cisTarget 2015 update: generalized cis-regulatory enrichment analysis in human, mouse and fly. Nucleic Acids Res., 43, W57–64.

46. Spletter, M.L., Barz, C., Yeroslaviz, A., Zhang, X., Lemke, S.B., Bonnard, A., Brunner, E., Cardone, G., Basler, K., Habermann, B.H., et al. (2018) A transcriptomics resource reveals a transcriptional transition during ordered sarcomere morphogenesis in flight muscle. Elife, 7, 1361.

47. Kaya-Çopur, A., Marchiano, F., Hein, M.Y., Alpern, D., Russeil, J., Luis, N.M., Mann, M., Deplancke, B., Habermann, B.H. and Schnorrer, F. (2021) The Hippo pathway controls myofibril assembly and muscle fiber growth by regulating sarcomeric gene expression. Elife, 10, 79.

48. Wilkinson, M.D., Dumontier, M., Aalbersberg, I.J.J., Appleton, G., Axton, M., Baak, A., Blomberg, N., Boiten, J.-W., da Silva Santos, L.B., Bourne, P.E., et al. (2016) The FAIR Guiding Principles for scientific data management and stewardship. Sci Data, 3, 160018–9.

49. Benjamini, Y. and Hochberg, Y. (1995) Controlling the False Discovery Rate: A Practical and Powerful Approach to Multiple Testing. Journal of the Royal Statistical Society: Series B (Methodological), 57, 289–300.

50. Xiao, Y., Hsiao, T.-H., Suresh, U., Chen, H.-I.H., Wu, X., Wolf, S.E. and Chen, Y. (2014) A novel significance score for gene selection and ranking. Bioinformatics, 30, 801–807.

51. Durinck, S., Moreau, Y., Kasprzyk, A., Davis, S., De Moor, B., Brazma, A. and Huber, W. (2005) BioMart and Bioconductor: a powerful link between biological databases and microarray data analysis. Bioinformatics, 21, 3439–3440.

52. Barrett, T., Wilhite, S.E., Ledoux, P., Evangelista, C., Kim, I.F., Tomashevsky, M., Marshall, K.A., Phillippy, K.H., Sherman, P.M., Holko, M., et al. (2013) NCBI GEO: archive for functional genomics data sets--update. Nucleic Acids Res., 41, D991–5.

53. Grau, J., Grosse, I. and Keilwagen, J. (2015) PRROC: computing and visualizing precision-recall and receiver operating characteristic curves in R. Bioinformatics, 31, 2595–2597.

54. Garcia-Alonso, L., Holland, C.H., Ibrahim, M.M., Turei, D. and Saez-Rodriguez, J. (2019) Benchmark and integration of resources for the estimation of human transcription factor activities. Genome Res., 29, 1363–1375.

55. Hardee, J., Ouyang, Z., Zhang, Y., Kundaje, A., Lacroute, P. and Snyder, M. (2013) STAT3 targets suggest mechanisms of aggressive tumorigenesis in diffuse large B-cell lymphoma. G3 (Bethesda), 3, 2173-2185.

56. Uslu, V.V., Petretich, M., Ruf, S., Langenfeld, K., Fonseca, N.A., Marioni, J.C. and Spitz, F. (2014) Long-range enhancers regulating Myc expression are required for normal facial morphogenesis. Nat. Genet., 46, 753–758.

57. Chen, A.T.-Y., Guo, C., Itani, O.A., Budaitis, B.G., Williams, T.W., Hopkins, C.E., McEachin, R.C., Pande, M., Grant, A.R., Yoshina, S., et al. (2015) Longevity Genes Revealed by Integrative Analysis of Isoform-Specific daf-16/FoxO Mutants of Caenorhabditis elegans. Genetics, 201, 613–629.

58. Schnorrer, F., Schönbauer, C., Langer, C.C.H., Dietzl, G., Novatchkova, M., Schernhuber, K., Fellner, M., Azaryan, A., Radolf, M., Stark, A., et al. (2010) Systematic genetic analysis of muscle morphogenesis and function in Drosophila. Nature, 464, 287–291.

59. Dietzl, G., Chen, D., Schnorrer, F., Su, K.-C., Barinova, Y., Fellner, M., Gasser, B., Kinsey, K., Oppel, S., Scheiblauer, S., et al. (2007) A genome-wide transgenic RNAi library for conditional gene inactivation in Drosophila. Nature, 448, 151–156.

60. Ni, J.-Q., Zhou, R., Czech, B., Liu, L.-P., Holderbaum, L., Yang-Zhou, D., Shim, H.-S., Tao, R., Handler, D., Karpowicz, P., et al. (2011) A genome-scale shRNA resource for transgenic RNAi in Drosophila. Nat. Methods, 8, 405–407.

61. Weitkunat, M. and Schnorrer, F. (2014) A guide to study Drosophila muscle biology. Methods, 68, 2–14.

62. Schindelin, J., Arganda-Carreras, I., Frise, E., Kaynig, V., Longair, M., Pietzsch, T., Preibisch, S., Rueden, C., Saalfeld, S., Schmid, B., et al. (2012) Fiji: an open-source platform for biological-image analysis. Nat. Methods, 9, 676-682.

63. Tepper, R.G., Ashraf, J., Kaletsky, R., Kleemann, G., Murphy, C.T. and Bussemaker, H.J. (2013) PQM-1 complements DAF-16 as a key transcriptional regulator of DAF-2-mediated development and longevity. Cell, 154, 676–690.

64. Wu, S., Liu, Y., Zheng, Y., Dong, J. and Pan, D. (2008) The TEAD/TEF family protein Scalloped mediates transcriptional output of the Hippo growth-regulatory pathway. Dev. Cell, 14, 388–398.

65. Wang, S., Reuveny, A. and Volk, T. (2015) Nesprin provides elastic properties to muscle nuclei by cooperating with spectraplakin and EB1. J. Cell Biol., 209, 529–538.

66. Kuleshov, M.V., Diaz, J.E.L., Flamholz, Z.N., Keenan, A.B., Lachmann, A., Wojciechowicz, M.L., Cagan, R.L. and Ma’ayan, A. (2019) modEnrichr: a suite of gene set enrichment analysis tools for model organisms. Nucleic Acids Res., 47, W183–W190.

67. Chèneby, J., Gheorghe, M., Artufel, M., Mathelier, A. and Ballester, B. (2018) ReMap 2018: an updated atlas of regulatory regions from an integrative analysis of DNA-binding ChIP-seq experiments. Nucleic Acids Res., 46, D267–D275.

68. Fuda, N.J., Guertin, M.J., Sharma, S., Danko, C.G., Martins, A.L., Siepel, A. and Lis, J.T. (2015) GAGA factor maintains nucleosome-free regions and has a role in RNA polymerase II recruitment to promoters. PLoS Genet, 11, e1005108.

